# CATSPERβ–δ Interaction Governs Hierarchical CatSper Holo-complex Assembly and is Essential for Male Fertility

**DOI:** 10.64898/2026.08.06.743317.1

**Authors:** Jong-Nam Oh, Huafeng Wang, Tse-en Wang, Xiaofang Huang, Leonard K. Kaczmarek, Jean-Ju Chung

## Abstract

The sperm-specific CatSper channel is macromolecular Ca2+ channel complex essential for hyperactivated motility and male fertility. The pore-forming channel (CATSPER1-4) associates with a large extracellular domain-containing canopy (CATSPERβ–ε), a cytosolic Ca²⁺ sensing subcomplex (CATSPERζ–EFCAB9–ARMH2), the putative transporter SLCO6C1, and CATSPERθ–η, arranging into higher-order zigzag rows in the flagellar membrane. However, the molecular mechanism governing holo-complex assembly during spermatogenesis remains largely undefined. Here we demonstrate that the CATSPERβ–δ interaction represents an essential early step in CatSper biogenesis. CRISPR/Cas9 targeting of *Catsperb* exon 4 generated a frameshift knockout (*Catsperb*^-/-^) and an in-frame deletion mutant (*Catsperb*^Δ/Δ^) that specifically disrupts the β–δ interaction interface. Disrupting this interface reduced CATSPERδ among canopy subunits, impairing canopy assembly and destabilizing the core pore-forming channel. Consequently, mature spermatozoa completely lacked the entire CatSper complex, phenocopying the knockout. A transgenic line expressing extracellular-domain-truncated CATSPERδ lacking the β-binding region likewise phenocopied the Catsperb mutant, confirming the necessity of an intact β–δ interface. AlphaFold-Multimer modeling and alanine-substitution mutagenesis identified key hydrogen-bonding residues at this interface that mediate canopy subunit association. Consistent with complex loss, whole-sperm patch-clamp recordings revealed complete absence of CatSper conductance in mutant spermatozoa. Both mutant lines exhibited defective sperm hyperactivation and male infertility despite normal spermatogenesis and baseline motility. Together, these findings establish that canopy formation driven by CATSPERβ–δ interaction precedes and is required for pore-forming channel assembly, defining a hierarchical assembly pathway for the CatSper holo-complex and highlighting a key structural node for male fertility and contraceptive development.

## Introduction

Multiprotein complexes are ubiquitous in both prokaryotes and eukaryotes, where more than half of all proteins function as dimers or higher-order assemblies; in vertebrates, approximately half of these complexes are heteromers [1, 2]. Among cellular proteins, membrane proteins occupy a critical position at the cell surface, mediating interactions between cells and sensing signals in their extracellular environment [3, 4]. The ordered assembly of multiprotein membrane complexes into functional units represents a fundamental yet poorly understood regulatory step in cell biology [5–8]. Hierarchical assembly pathways are well illustrated by voltage-gated K^+^ channels, where the N-terminal cytosolic Tetramerization (T1) domain drives subfamily-specific recognition to prevent mismatched cross-talk between distinct lineages (e.g., *Shaker* vs. *Shab*) [9, 10]. This core tetramer subsequently establishes a four-fold symmetric platform [11] that docks K_v_β subunits to form a functional macromolecular octet (4α: 4β) [12]. While the T1 domain is a paradigm for four-fold (C4) symmetry, evolutionary adaptations of this structural module display remarkable architectural plasticity. Notably, the soluble K^+^ channel tetramerization domain-containing (KCTD) family of proteins leverages a homologous BTB-like fold to drive homopentameric (C5) configurations instead of tetramers, expanding the functional repertoire of the scaffold into diverse signaling networks [13, 14].

While these classical somatic models provide elegant paradigms for symmetrical oligomerization, highly specialized cells like spermatozoa break these structural conventions, imposing far more complex demands on macromolecular coordination to navigate their environment [15–18]. Achieving the precise physiological responses required for fertilization hinges on an unprecedented level of protein biogenesis and spatial organization within the flagellar membrane. At the center of this challenge is CatSper, the sperm-specific, voltage-dependent, and alkalinization activated Ca²⁺ channel that is absolutely required for male fertility, serving as the primary Ca²⁺ entry pathway that drives hyperactivated motility — the vigorous, asymmetric flagellar beating required for successful fertilization [19, 20]. In stark contrast to simple tetramers, CatSper is a staggering, multi-layered asymmetric heteromeric complex composed of at least 15 components [21–23]. An asymmetric pore-forming core comprised of four distinct α-subunits (CATSPER1, 2, 3, 4) [24–27] is associated with four distinct single-transmembrane (TM) auxiliary subunits bearing large extracellular domains (ECDs) — CATSPERγ, ε, δ, β, which assemble a canopy-like structure in a defined arrangement above the pore [28–31]. The channel further associates with a cytoplasmic Ca^2+^-sensing sub-complex formed by EFCAB9, CATSPERζ, and ARMH2 [21, 32–34]. Two of these holo-complexes are rotated by 180° to form a dimer, which is linked extracellularly by the β-β interface and at the transmembrane region by two additional small TM subunits (CATSPERθ, η) [22, 23, 35, 36]. In addition, an organic anion transporter, SLCO6C1, associates with each holo-complex at the periphery of the dimers, and dimers assemble into quadrilinear zigzag rows along the flagellar membrane [22, 23, 36], forming structurally distinct Ca²⁺-signaling nanodomains [37]. Genetic ablation of any TM subunit in mice results in loss of the entire CatSper complex from mature spermatozoa [19, 20, 29, 30, 35], a phenotypic outcome mirrored in infertile human subjects [38–42]. While these findings underscore a strict, obligate interdependence of subunits for complex stability of the CatSper channel in mice, how individual components are selectively recruited and incorporated into this mature holo-complex during spermatogenesis is only beginning to be defined [29, 30].

Our previous genetic studies showed that loss of either CATSPERδ or CATSPERε compromises the stability of CATSPER1 in the testis [29, 30], whereas CATSPER4 level is reduced in the absence of CATSPERθ [35], indicating that auxiliary subunits play indispensable roles in channel biogenesis. Notably, transcripts encoding these single-TM auxiliary subunits are expressed prior to the pore-forming α-subunits [29, 30]. This suggests a stepwise, hierarchically regulated assembly process wherein the pre-formed canopy serves as a structural scaffold to recruit and stabilize the core α-subunit tetramer [30]. Here, we build on this model by focusing on a CRISPR/Cas9-induced spliced region in CATSPERβ that contributes to a protein-protein interaction (PPI) interface with CATSPERδ. We generated a mouse line carrying a targeted in-frame mutation in this region of CATSPERβ that disrupts CATSPER β–δ interaction, thereby perturbing a key node in the proposed assembly pathway. Strikingly, the CATSPERβ mutant exhibited loss of the entire CatSper complex in spermatozoa, accompanied by absence of hyperactivated motility and male fertility. Transgenic expression of an ECD-truncated CATSPERδ lacking the β–δ binding region phenocopied the CATSPERβ mutant. Finally, alanine substitution of predicted interface residues in CATSPERβ reduced β– δ binding affinity. Together, these results identify the CATSPERβ–δ interaction as a critical determinant of canopy formation and provide mechanistic support for a hierarchical order of CatSper biogenesis during spermatogenesis.

## Material and Methods

### Animals

Wild-type C57BL/6J mice were purchased from the Jackson Laboratory. *Catsper1-null* (RRID:MGI:3694476) and *Catsperd-null* (RRID:MGI: 2147030) mice [19, 29] were previously generated and maintained on a C57BL/6 background. All animal experiments were conducted in accordance with guidelines approved by the Yale Institutional Animal Care and Use Committee (IACUC, #20079).

### Antibodies and Reagents

Rabbit polyclonal antibodies specific to mouse CATSPER1, CATSPER3, CATSPER4, CATSPERδ, CATSPERγ, CATSPERε and EFCAB9 were described previously [19, 21, 27, 29, 30, 32]. To generate a rabbit polyclonal antibody against mouse CATSPERβ, a peptide corresponding to residues 169–186 of mouse CATSPERβ (EMEVRDLYPEVNDIKVTK) was synthesized and conjugated to KLH carrier protein (Open Biosystems). Antisera from immunized rabbits were affinity-purified using the peptide immobilized on AminoLink Plus resin (Pierce, Thermo Scientific). Anti-acetylated α-tubulin antibody (clone 6-11B-1; Sigma-Aldrich, T7451), anti-Myc antibody (Sigma-Aldrich, OP10), anti-Flag antibody (Cell signaling, 8146), anti-HA antibody (Cell signaling, 2367), anti-beta Actin antibody (Cell signaling, 4970), anti-Phosphotyrosine Antibody (4G10, Sigma-Aldrich, 05-1050X), and anti-mCherry antibody (Novus Biologicals, NBP1-96752) were purchased commercially. Unless otherwise indicated, all chemicals were obtained from Sigma-Aldrich.

### Generation of *Catsperb*-Knockout and Internal Deletion Mutant Mouse Lines and Genotyping

*Catsperb*-null (*Catsperb*^−/−^) and in-frame deletion mutant (*Catsperb*^Δ/Δ^) mouse lines were generated on a C57BL/6J background using the CRISPR/Cas9 system. Superovulated female mice were mated with wild-type males to obtain fertilized eggs. A guide RNA (5’-ACGAACAGTACATTCTACGG-3’) targeting the fourth exon of mouse *Catsperb* was microinjected into the pronuclei of fertilized eggs, and the resulting two-cell stage embryos were transferred into the oviducts of pseudopregnant recipient females. The target region was amplified by polymerase chain reaction (PCR) from alkaline-lysed tail biopsies of founder pups using the following primers: forward, 5’-AAAGTCCGCTGTTTCTTCAGAA-3’; reverse, 5’-GAGTCACACTTGTACAACCAC-3’. The resulting PCR fragments were subjected to Sanger sequencing to confirm genome editing at the target site. Monoallelic founders carrying 10 bp and 11 bp insertions at the target site were backcrossed to wild-type C57BL/6J animals to confirm germline transmission of the mutant alleles. Routine genotyping of offspring was performed using wild-type *Catsperb*-specific primers (forward, 5’-AACAGTACATTCTACGGTGCTA-3’; reverse, 5’-AAGGCATCCAATAAGAAAGAGTGTG-3’).

### Generation of *Catsperd*-Tg mice expressing ECD-truncated CATSPERδ

The generation of transgenic mice was conducted according to the reported protocol [30]. In brief, the transgene encoding the ECD-truncated CATSPERδ (*Catsperd*-TG) was introduced into the pClgn-mCherry vector, and the linearized plasmid was then electroporated into fertilized eggs collected from C57BL/6 mice (**Supplemental Fig.** 7). The extracted gDNAs of the founders were subjected to PCR using the following primers: forward primer, 5′-AGTTGCCCAGAAACATCCAG-3′; and reverse primer, 5′-TGAAGCGCATGAACTCCTT-3′. *Catsperd*-null;*Catsperd*-TG mice were generated by crossing *Catsperd*-TG males with *Catsperd*-null females.

### Mating Test

To assess male fertility, wild-type, *Catsperb*^+/−^, *Catsperb*^−/−^, *Catsperb*^Δ/Δ^, *Catsperd*^+/−;TG+^ and *Catsperd*^−/−;TG+^ males were each individually housed with a fertile wild-type female and monitored over a two-month period. Pregnancy rate and litter size were recorded for each mating pair.

### Preparation of Mouse Spermatozoa

Epididymal spermatozoa from adult male mice (8–16 weeks of age) were collected by swim-out from cauda epididymis into M2 medium (Millipore). For capacitation, collected spermatozoa were transferred to human tubular fluid (HTF) medium (Millipore) at a concentration of 2 × 10⁶ cells/ml and incubated for 90 min at 37°C under 5% CO₂.

### Sperm Motility Analysis

#### Computer-assisted sperm analysis

Computer-assisted sperm analysis (CASA) was performed as previously described [33]. Aliquots of non-capacitated and capacitated spermatozoa (2 × 10⁶ cells/ml) were loaded into CellVision slide chambers and analyzed on a 37°C heated stage of a Nikon E200 microscope equipped with a 10X phase-contrast objective (CFI Plan Achro 10X/0.25 Ph1 BM, Nikon). Images were captured at 50 fps using a CMOS camera (Basler acA1300-200um; Basler AG, Ahrensburg, Germany) and analyzed using Sperm Class Analyzer software (v6.6; Microptic, Barcelona, Spain). A minimum of 200 motile spermatozoa were analyzed per trial, with at least three biological replicates performed per genotype.

#### Flagellar Waveform Analysis

To tether sperm heads for planar flagellar beating, non-capacitated and capacitated spermatozoa (2 × 10⁶ cells/ml) from adult mice were transferred to 37°C HEPES-buffered HTF medium [32] in fibronectin-coated Delta T chambers (Bioptechs). Flagellar movements of tethered spermatozoa were recorded for 2 s at 200 frames per second (fps) using a pco.edge sCMOS camera mounted on an Axio Observer Z1 microscope (Carl Zeiss). All recordings were completed within 10 min of transferring spermatozoa to the imaging chamber. Recorded image stacks were processed using FIJI [43] to measure beat frequency and α-angle of the sperm flagellum [29], and to generate overlaid images tracing flagellar waveforms, as previously described [32].

#### Free-Swimming in High Viscosity Assay

Prior to and following capacitation, spermatozoa were transferred to 37°C HEPES-buffered HTF medium supplemented with 0.3% methylcellulose and placed in Delta T chambers. Sperm movement was recorded, and superimposed trajectory images were generated from the video recordings using the same imaging and analysis pipeline described for the flagellar waveform analysis [44].

### Electrophysiological Recording

Corpus epididymal spermatozoa were released into and washed with HEPES-buffered saline and subsequently attached onto a 35-mm culture dish. Gigaohm seals were formed at the cytoplasmic droplet of motile spermatozoa as previously described [45]. Whole-cell CatSper currents were recorded from wild-type, *Catsperb*^Δ/Δ^, and *Catsperb*^−/−^ spermatozoa in a divalent-free bath solution containing (in mM): 150 Na-gluconate, 20 HEPES, and 5 Na₃HEDTA, pH 7.4. The intrapipette solution consisted of (in mM): 135 Cs-methanesulfonate (CsMes), 10 HEPES, 10 EGTA (ethylene glycol tetraacetic acid), and 5 CsCl, adjusted to pH 7.2 with CsOH. Data were sampled at 10 kHz and low-pass filtered at 2 kHz. Current recordings were analyzed using Clampfit software (Axon Instruments, Gilze, Netherlands), and Figures were plotted using Grapher 8 (Golden Software, Inc., Golden, CO).

### Quantitative Reverse Transcription PCR

Total RNA was purified from testes of wild-type, *Catsperb*^−/−^, and *Catsperb*^Δ/Δ^ mice using the RNeasy Mini Kit (QIAGEN, Netherlands) according to the manufacturer’s instructions. Five hundred nanograms of purified RNA was reverse-transcribed using the iScript cDNA Synthesis Kit (Bio-Rad). The resulting cDNAs were subjected to quantitative PCR (qPCR) on a CFX96 real-time system (Bio-Rad) using SYBR Green qPCR mix (ABclonal, MA). To identify splicing of mutant transcripts, primer pairs spanning the CRISPR/Cas9 targeting site within exon 4 of *Catsperb* were used; transcript levels were further assessed using primers spanning exon– exon junctions (exon 1–2, exon 6–7, and exon 7–8). The presence of wild-type and transgenic *Catsperd* was verified by quantitative PCR. *Gapdh* and *Tbp* were used as reference genes for normalization. All primer sequences are listed in Table 1.

**Table 1.**
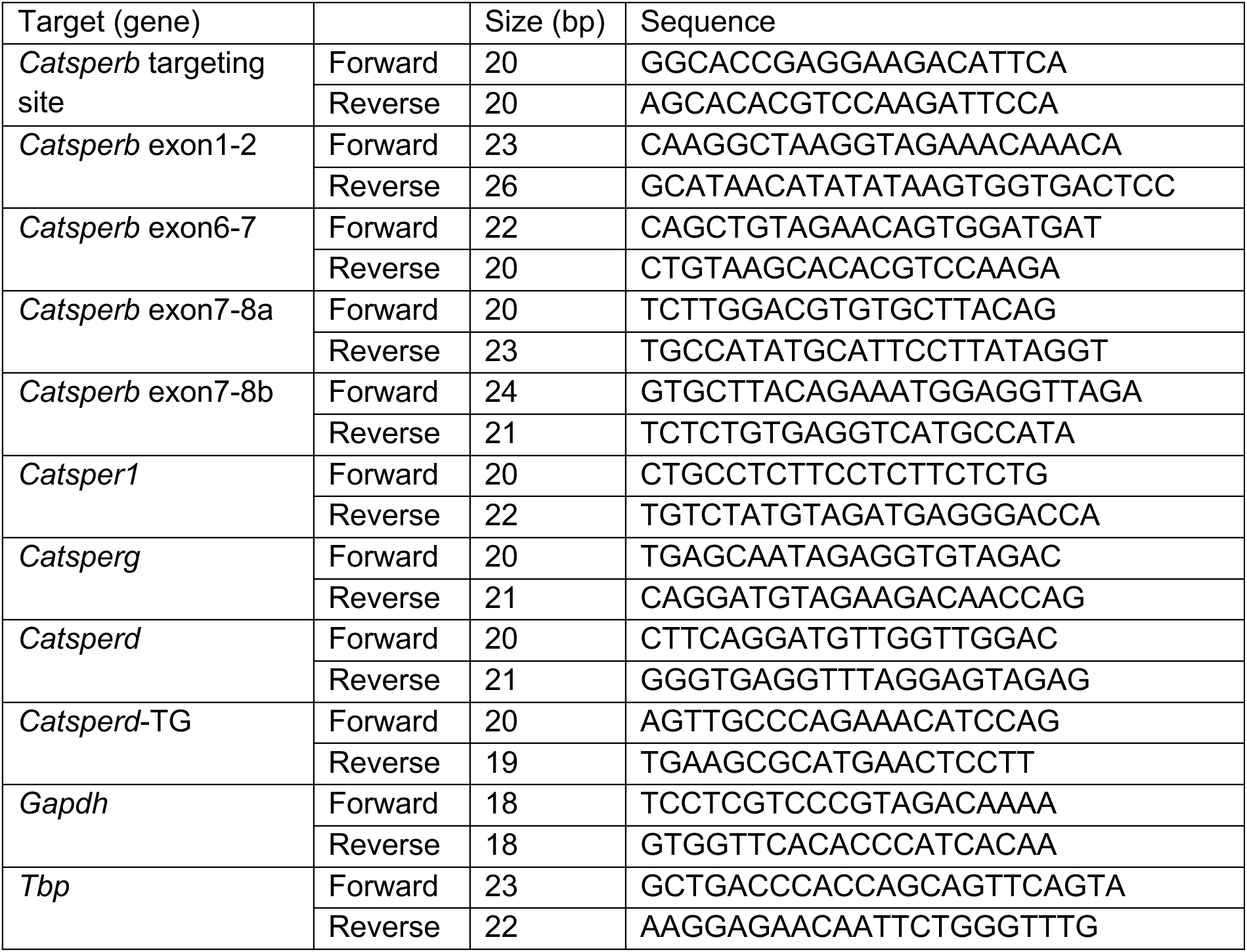
Primers used for quantitative RT-PCR.

### Protein Extraction and Immunoblotting of Epididymal Spermatozoa

Epididymal spermatozoa washed in phosphate-buffered saline (PBS) were directly lysed in 2X LDS sample buffer for 10 min with vortex mixing at room temperature (RT). Lysates were cleared by centrifugation at 18,000 × g for 10 min at 4°C, and the resulting supernatants were denatured by the addition of DTT to a final concentration of 50 mM and incubation at 75°C for 10 min. Denatured proteins were resolved by SDS-PAGE and transferred for immunoblotting. Primary antibodies used are: rabbit polyclonal anti-mouse CATSPERβ, CATSPERδ, CATSPER1, CATSPER3, and EFCAB9 (each at 1 µg/ml), and anti-acetylated α-tubulin (AcTUB; 50 ng/ml). HRP-conjugated goat anti-rabbit or anti-mouse secondary antibodies (Jackson ImmunoResearch) were used at 1:20,000 dilution according to the host species of the primary antibody.

### Protein Extraction, Immunoblotting, and Co-Immunoprecipitation of Testicular Proteins

Mouse testes were homogenized in 320 mM sucrose (1:10, wt/vol), and cell debris and nuclei were removed by centrifugation at 1,000 × g for 20 min at 4°C. The post-nuclear supernatant was subjected to ultracentrifugation at 105,000 × g for 1 hr at 4°C to collect the microsomal membrane fraction. The microsomal pellet was solubilized in PBS containing 1% Triton X-100 and protease inhibitor cocktail (cOmplete Mini, Roche) by rocking at 4°C for 3 hr. The detergent-insoluble material was removed by centrifugation at 18,000 × g for 1 hr at 4°C. The clarified supernatant (solubilized microsome) was used directly for SDS-PAGE and immunoblotting or for co-immunoprecipitation. For co-immunoprecipitation, solubilized microsomal proteins were incubated overnight at 4°C with SureBeads Protein A Magnetic Beads (Bio-Rad) pre-conjugated with 0.5 µg of anti-CATSPERδ or anti-CATSPERγ antibody. For immunoblot detection following co-immunoprecipitation, Clean-Blot IP Detection Reagent (Thermo Scientific) and VeriBlot for IP Detection Reagent (Abcam) were used at 1:200 dilution to minimize interference from immunoglobulin heavy and light chains.

### Immunocytochemistry

Epididymal spermatozoa were washed in PBS and attached onto glass coverslips by centrifugation at 700 × g for 5 min. Attached cells were fixed with either 100% methanol or 4% paraformaldehyde (PFA) for 10 min at RT, and washed three times with PBS. Fixed cells were permeabilized with 0.1% Triton X-100 in PBS for 10 min and subsequently blocked with 10% normal goat serum in PBS for 1 hr at RT. Blocked samples were incubated with primary antibodies against CATSPER1 and CATSPERβ (each at 10 µg/ml) overnight at 4°C. Alexa Fluor 568-conjugated goat anti-rabbit secondary antibody (0.1 µg/ml; Invitrogen) was applied for 1 hr at RT. Coverslips were mounted with ProLong Gold Antifade Reagent with DAPI (Invitrogen). Images were acquired by confocal microscopy using a Zeiss LSM710 Elyra P1 (software: Zen 2012 SP2(black)) system equipped with a Plan-Apochromat 63×/1.40 oil immersion objective (Carl Zeiss).

### Co-expression of CATSPERδ and Mutant CATSPERβ in a Heterologous System

To assess the functional significance of amino acid residues identified at the CATSPERβ–δ interface by structural prediction, plasmids encoding wild-type or alanine-substituted mutant CATSPERβ-FLAG were co-transfected with CATSPERδ-Myc into HEK293T cells (**Fig. 6A**). At 48 h post-transfection, cells were lysed and subjected to anti-Myc immunoprecipitation, followed by immunoblotting to detect co-precipitated FLAG-tagged CATSPERβ proteins.

### AlphaFold Structure Prediction and Structural Analysis

The structures of mutant CATSPERβ proteins were predicted using AlphaFold2 [46], and dimer structures of CatSper subunit pairs were predicted using AlphaFold-Multimer, both implemented via Google Colab notebooks [47]. The protein sequences used as inputs for structure prediction are listed in Table S1. Predicted structures were visualized and analyzed using UCSF ChimeraX [48]. Predicted models were fitted onto the atomic structure of the CatSper channel complex (PDB: 7EEB) using the *matchmaker* tool in ChimeraX to determine the location of predicted structures within the holo-complex. Hydrogen bond contacts between chains were identified using a distance cutoff of 2 Å with tolerances of 0.1, 0.5, and 1 Å.

### Quantification and Statistical Analysis

All experiments were performed at least three times. Sample sizes for each experiment are indicated in the corresponding Figure legends. Graphs were plotted using Prism 11 (GraphPad, MA). Statistical analyses were performed using Student’s *t*-test or one-way ANOVA as appropriate. Data are presented as mean ± SEM. Differences were considered statistically significant at *p<0.05, **p<0.01, ***p<0.001, and ****p<0.0001.

## Results

### Strategic CRISPR/Cas9 targeting of *Catsperb* exon 4 induced aberrant splicing, simultaneously generating both internal deletion-mutant and knockout mouse lines

The genes encoding CatSper canopy subunits (β, γ, δ, and ε) are expressed earlier than those encoding the pore-forming subunits (CATSPER1–4) during spermatogenesis (GSE109033) [29, 32, 49], suggesting that the canopy assembles likely precedes pore-formation. At the structural level, CATSPERβ anchors the dimeric interface within the transmembrane domain and extends outward to interact with the neighboring δ and γ subunits within the extracellular canopy (**Fig. 1A-B**) [22, 23, 36]. This distinctive structural position suggests that CATSPERβ is critical not only for assembling the higher-order supramolecular CatSper organization along the flagellum, but also for assembling the individual holo-complex. We therefore hypothesize that the CATSPERβ-mediated protein-protein interactions (PPIs) at the canopy level drive subsequent CatSper assembly and functional regulation.

**Figure 1.**
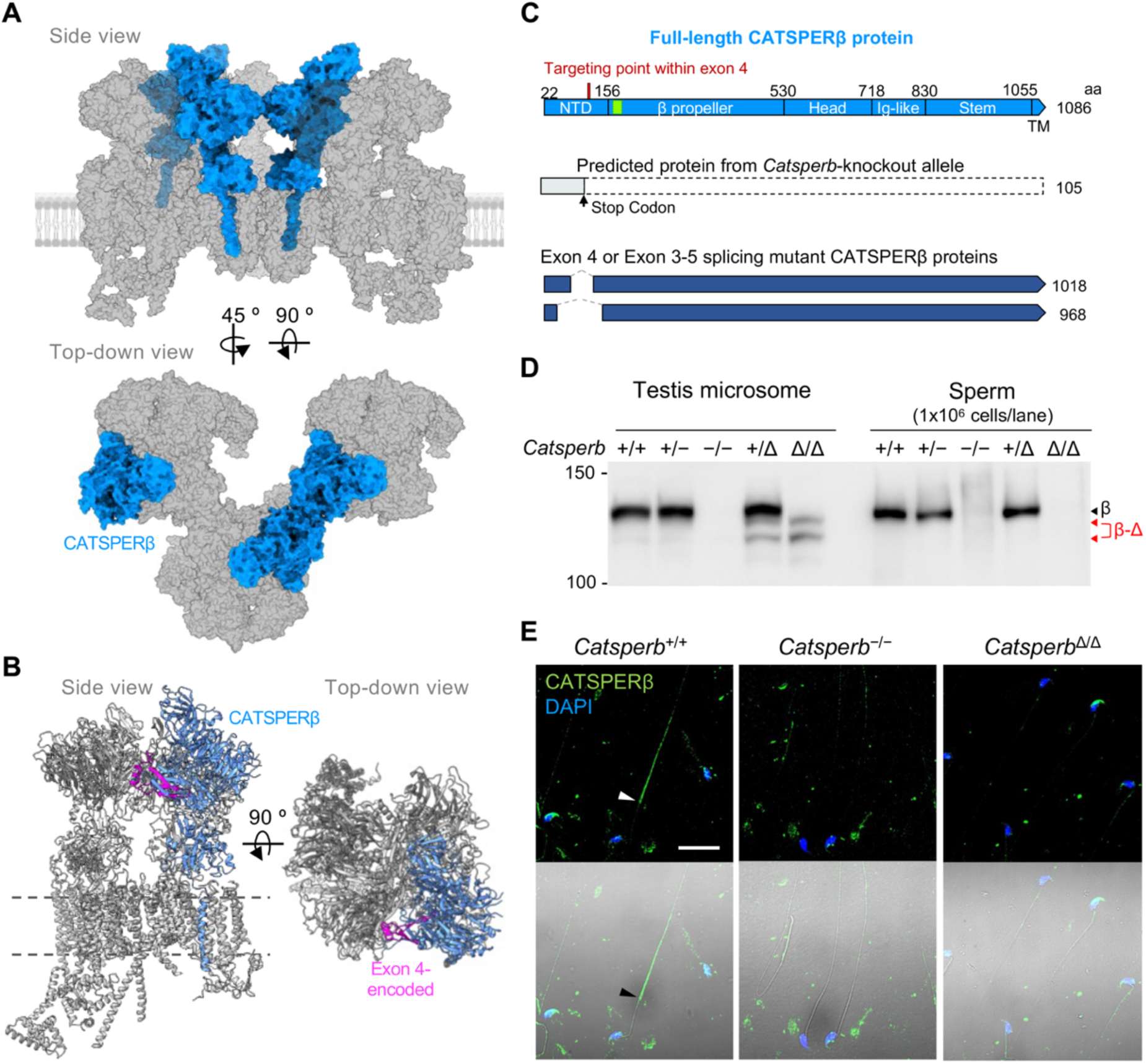
Generation of *Catsperb*-knockout and internal deletion mutant mouse models by CRISPR/Cas9 system. (A) Surface display of mouse CatSper trimer (PDB: 9UBX). CATSPERβ is shown in blue. (B) Atomic model of the CatSper channel complex (PDB: 7EEB) with CATSPERβ and its exon 4-encoded region are shown in blue and magenta, respectively. (C) Schematic diagram of CATSPERβ domain organization and predicted mutant CATSPERβ proteins in the genome-edited mouse models. The red mark shows the targeting point of CRISPR/Cas9 in exon 4. The marked green box indicates the epitope location for anti-CATSPERβ antibody used in the study (α-CATSPERβ (169), *top*). CRISPR/Cas9 introduced frameshift indels and exon skipping to establish Catsperb-knockout and internal deletion mutant mice, respectively, which are predicted to encode two internally deleted CATSPERβ proteins. (D) Detection of full-length and internally deleted CATSPERβ proteins by immunoblot analysis in testis microsomes and whole sperm lysates from wild type (+/+), *Catsperb* knockout (+/− and −/−) and mutant (Δ/+ and Δ/Δ) mice. (E) Detection of CATSPERβ in the principal piece of sperm flagella by immunocytochemistry in the spermatozoa from wild-type but not in those from Catsperb knockout and homozygous mutant animals. *See also* Supplemental Figure 1.

To test this idea, we used CRISPR/Cas9 to specifically target *Catsperb* exon 4, which encodes the CATSPERδ-interfacing region, aiming to obtain both an in-frame deletion mutant (*Catsperb*^Δ/Δ^) and an early-termination frameshift knockout (*Catsperb*^−/−^) (**Fig. 1C**; **Supplemental Fig. 1**). Founder screening identified one allele introducing a premature stop codon within the CATSPERβ N-terminal β-propeller domain and another allele that induced exon skipping (exon 4 alone or exons 3–5 together), thereby simultaneously establishing the knockout and internal deletion mutant lines, respectively (**Supplemental Fig. 1**).

*Catsperb* transcript levels were significantly reduced in *Catsperb*^−/−^ males but were unchanged in *Catsperb*^Δ/Δ^ males relative to wild type (**Fig. 2A-B**; **Supplemental Fig. 2A**), indicating that exon skipping did not impair transcription or mutant transcript stability. Consistently, internally deleted CATSPERβ protein was detected in the testes of *Catsperb*^Δ/Δ^ males, whereas full-length CATSPERβ was absent in *Catsperb*^−/−^ testes (**Fig. 1D**; **Fig. 2C, E**). However, epididymal spermatozoa from both *Catsperb*^Δ/Δ^ and *Catsperb*^−/−^ males lacked CATSPERβ by Western blot and immunocytochemistry (**Fig. 1D-E**). These results show that truncating the CATSPERδ-interfacing CATSPERβ ECD is sufficient to cause complete loss of the CatSper complex from mature spermatozoa.

**Figure 2.**
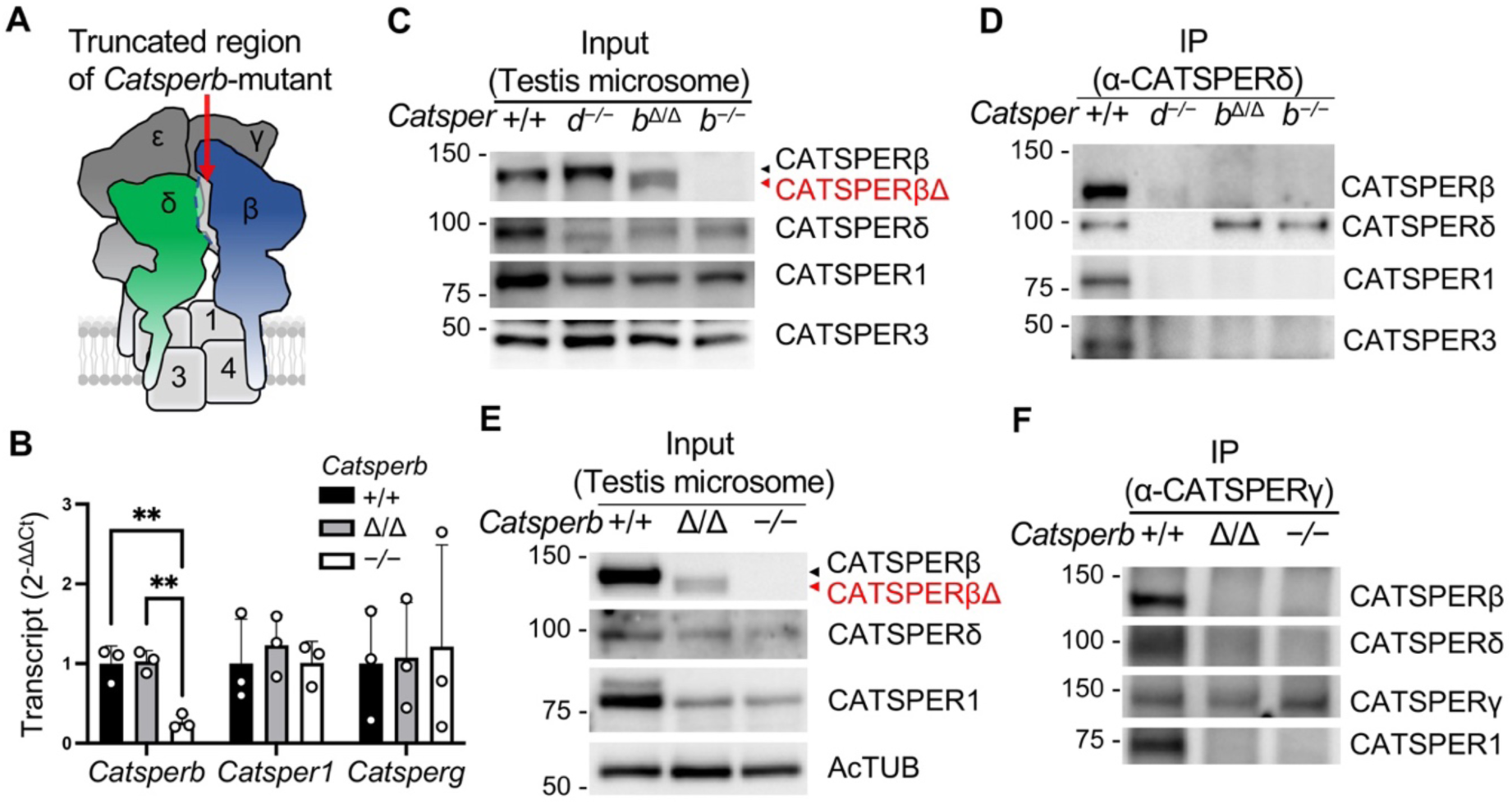
Internal deletion in the CATSPERβ NTD impairs the assembly of the channel complex, as does by complete loss of CATSPERβ. (A) Cartoons depicting the predicted channel complex during channel assembly in *Catsperb* in-frame mutant and knockout animals. (B) Relative levels of CatSper transcripts (*Catsperb*, *Catsper1*, *Catsperg*) in adult testis. (C and E) Protein levels of CatSper subunits in testis microsome from wildtype, *Catsperd*^−/−^, *Catsperb*^Δ/Δ^, and *Catsperb*^−/−^ by immunoblot analysis. Input for Figure 2D and 2F, respectively. Red arrow indicates internally deleted CATSPERβ protein (Δβ). (D and F) Detection of CatSper proteins in anti-CATSPERδ (D) and anti-CATSPERγ immunocomplex (F). *See also* Supplemental Figure 2, 3, and 4.

### Disrupting the CATSPERβ–δ interface deregulates CatSper canopy assembly

To assess how internal deletion of CATSPERβ at the β–δ interface affects CatSper complex assembly, we fitted AlphaFold-predicted structures of the CATSPERβ mutant into the CatSper atomic model (PDB: 7EEB). The predicted fold was overall similar to the full-length protein (**Fig. 2A**; **Supplemental Fig. 2B-C**) but the internally deleted mutants are predicted to reduce the extent of CATSPERβ-δ binding (**Supplemental Fig. 2D-F**). We then tested whether CATSPERδ remains associated with the CATSPERβ mutants by co-immunoprecipitation (Co-IP). Although *Catsperb* transcript levels were unchanged and CATSPERβ mutant proteins were detectable in *Catsperb*^Δ/Δ^ testis (**Fig. 1D**; **Fig. 2B, C, E**), the mutant CATSPERβ was not recovered in the CATSPERδ immunocomplex, mirroring the result observed in *Catsperb*^−/−^ and *Catsperd*^−/−^ testes (**Fig. 2D**).

Notably, the protein level of CATSPERδ, but not the other canopy subunits γ and ε, was also reduced in both *Catsperb*^Δ/Δ^ and *Catsperb*^−/−^ testes (**Fig. 2C, E**; **Supplemental Fig. 2G-I**; *see also* **Fig. 5B**; **Supplemental Fig. 7E-F**). Conversely, the absence of CATSPERδ [29] or the expression of canopy roof-truncated CATSPERδ in *Catsperd-/-* background (*Catsperd*^−/−;TG+^) did not affect the CATSPERβ and CATSPERγ protein levels in testes (*see* **Fig. 5B**; **Supplemental Fig. 7E**). Additionally, the internally deleted CATSPERβ was also absent from the CATSPERγ immunocomplex in *Catsperb*^Δ/Δ^ testis (**Fig. 2F**), even though the deleted region is not predicted to directly mediate CATSPERβ-CATSPERγ interactions. In comparison, CATSPERε—a direct interacting partner of CATSPERγ but not in contact CATSPERβ— remained detectable in the CATSPERγ immunocomplex in *Catsperb*^Δ/Δ^ testes (*see* **Supplemental Fig. 7E-F**). Together, these results support that the in-frame deletion does not measurably alter mRNA expression of other *CatSper* genes (i.e., *Catsper1* and *Catsperg*) (**Fig. 2B**), but that the expressed CATSPERβ mutant that disrupting CATSPERβ–δ interface deregulates complete CatSper canopy assembly.

### The mutant CATSPERβ partly loses its interface with CATSPERδ

To evaluate the structural impact of *Catsperb* exon skipping, we used AlphaFold-Multimer [50] to model canopy subunit interfaces using wild-type or mutant CATSPERβ together with CATSPERδ and CATSPERγ (ECD-only). The predicted dimers fit well onto the CatSper canopy structure (**Supplemental Fig. 3**; **Movie S1**). Both wild-type and internal deletion mutant CATSPERβ are predicted to engage CATSPERδ in a manner broadly consistent with the published CatSper complex structure (**Supplemental Fig. 3B-C; Supplemental Fig. 4A-B**; **Movies S2-S3**). Likewise, the CATSPERβ-CATSPERγ interfaces were comparable between wild-type and mutant, with no discernible change in β–γ binding (**Supplemental Fig. 4C-D**; **Movie S2**). Notably, internal deletion of the exon 4-encoded 49 amino acids within the β–δ interface reduced intermolecular proximity in the mutant (**Supplemental Fig. 4E-F**; **Movies S3-4**). Thus, while the mutant CATSPERβ retains part of the CATSPERδ-binding surface, the β–δ interaction is likely structurally compromised.

### Incomplete canopy formation impairs stability of the tetrameric CatSper pore-forming channel

Next, we tested stability of the pore-forming subunits in *Catsperb*^−/−^ and *Catsperb*^Δ/Δ^ testes by performing immunoblotting for CATSPER1, CATSPER3, and CATSPER4. Consistent with prior reports in *Catsperd*-and *Catspere*-knockout mice [15, 30], CATSPER1 protein levels, as well as CATSPER3 and CATSPER4, were reduced in the testes of both *Catsperb* mutants relative to wild type (**Fig. 2C, E**; **Supplemental Fig. 2J**; *see* **Supplemental Fig. 7E**). These data corroborate that the pre-assembly of the canopy structure, driven by CATSPERβ–δ interaction, is a prerequisite for formation of the tetrameric pore-forming channel and the holo-complex assembly.

### CATSPERβ internal deletion mutant and knockout spermatozoa similarly lack functional CatSper channels, leading to defective sperm hyperactivation and male infertility

Because canopy assembly is required for pore formation, we predicted that functional CatSper would be absent from spermatozoa of both *Catsperb*^Δ/Δ^ and *Catsperb*^−/−^ males. Consistent with this prediction, immunoblot analysis and immunocytochemistry revealed that, in spermatozoa from both genotypes, not only CATSPERβ but also all other ancillary and pore-forming subunits examined were undetectable (**Fig. 3A-B**). Whole-cell patch-clamp recordings corroborated these biochemical findings (**Fig. 3C-E**). Under divalent-free (DVF) conditions, monovalent *I_CatSper_* was not detected in either mutant or knockout spermatozoa using step or ramp voltage protocols (**Fig. 3C-D**). In contrast, wild-type spermatozoa exhibited robust *I_CatSper_* (**Fig. 3D-E**), the subsequent perfusion with HS solution containing calcium served to confirm the identity of the recorded current, validating the integrity of our electrophysiological approach.

**Figure 3.**
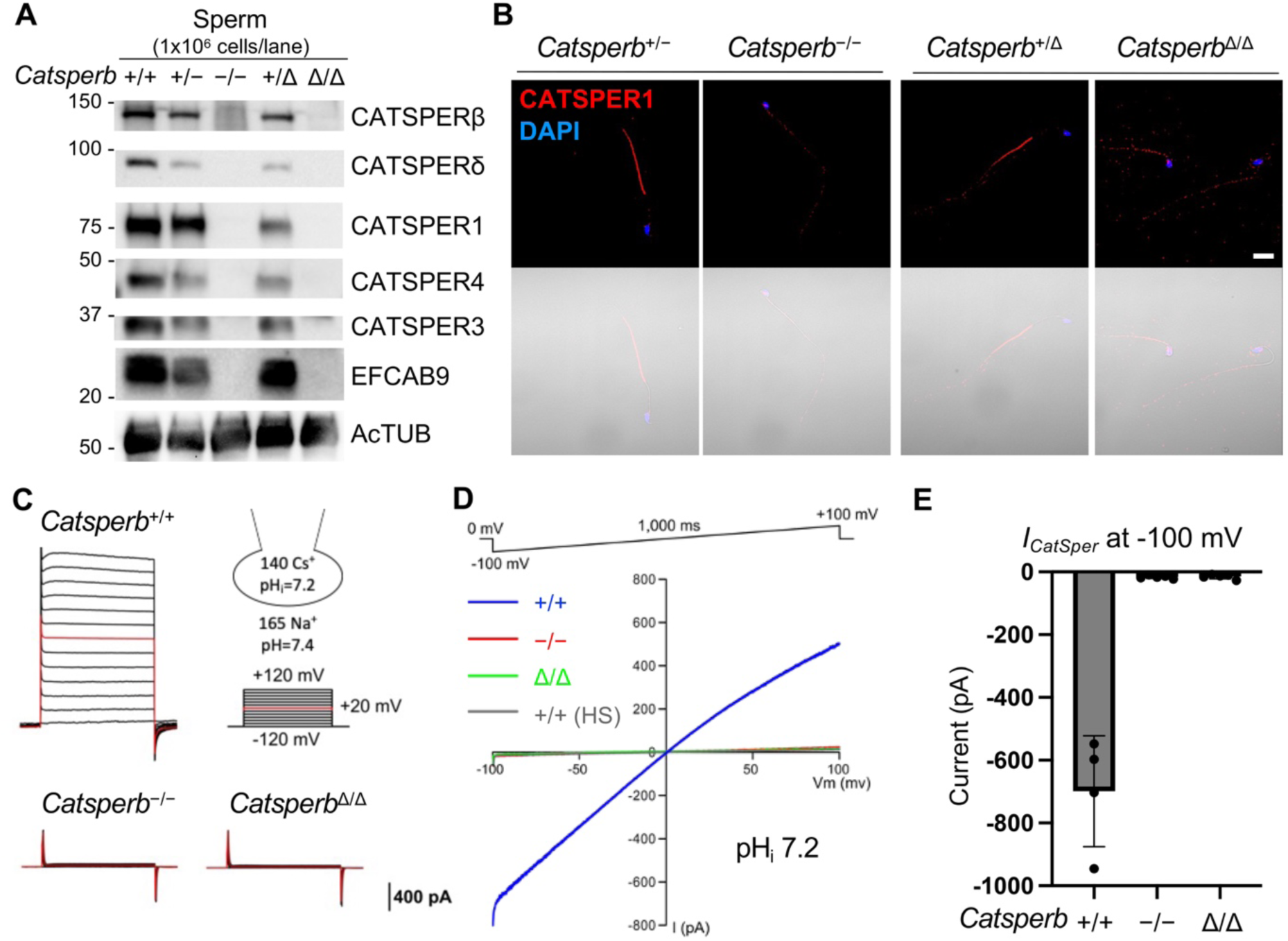
Functional CatSper channels are not assembled in the sperm from both Catsperb-knockout and in-frame deletion mutant mice. (A and B) Absence of the entire CatSper channel, shown by immunoblot analysis of CatSper components (A) and immunocytochemistry with anti-CATSPER1 antibody (B) of *Catsperb*-knockout and –mutant sperm. (C and D) Representative traces of CatSper current (*I_CatSper_*) from WT, *Catsperb*^−/−^, and *Catsperb*^Δ/Δ^ spermatozoa. *I_CatSper_* was elicited by a step protocol (–120 mV to +120 mV) in 20 mV increments (C) or by a voltage ramp (100 mV to +100 mV) from 0 mV holding potential (D). (E) Inward *I_CatSper_* at –100 mV from WT (N = 4), *Catsperb*^−/−^ (N = 5), and *Catsperb*^Δ/Δ^ (N = 5) spermatozoa.

Consistent with the biochemical and electrophysiological results, male fertility and sperm hyperactivation were severely impaired in both *Catsperb*^Δ/Δ^ and *Catsperb*^−/−^ mice (**Fig. 4A-D**), despite normal sperm count and total motility (**Supplemental Fig. 5**), matching phenotypes reported for other knockouts of *CatSper* genes encoding TM components [29, 30]. After 90 min incubation under capacitating conditions, both mutant and knockout spermatozoa exhibited reduced flagellar beat amplitude and loss of waveform asymmetry (**Fig. 4E-F**; **Supplemental Fig. 6**; **Movie S5**), together with decreased beat frequency and an increased α-angle after capacitation (**Fig. 4G-J**), features associated with sperm hyperactivation.

**Figure 4.**
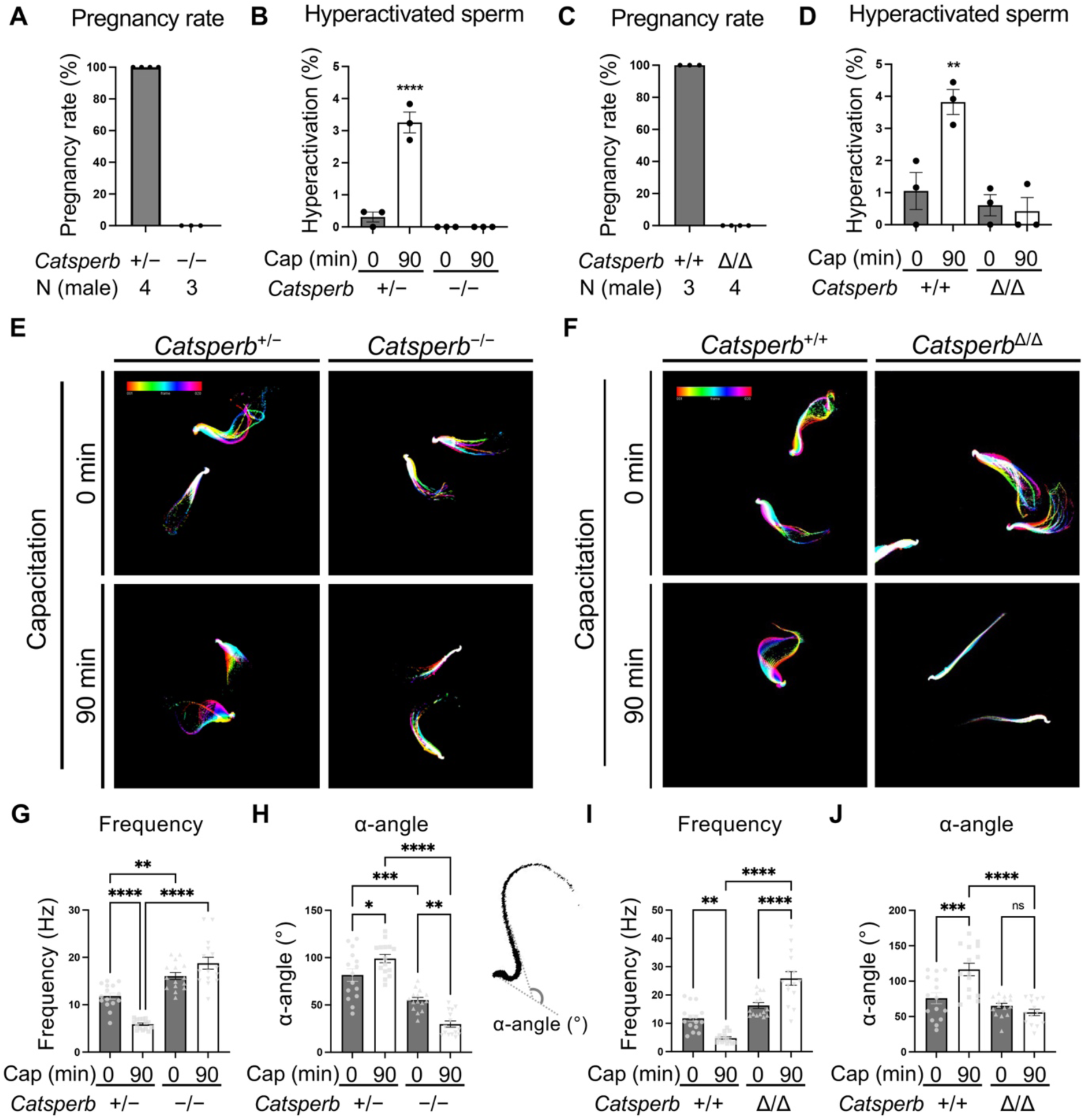
Sperm hyperactivation is impaired in both *Catsperb*-knockout and in-frame deletion mutant males. (A and C) Pregnancy rates of fertile females mated with *Catsperb*^−/−^ (A) or *Catsperb*^Δ/Δ^ (C) males compared to their littermate *Catsperb*^+/−^ or *Catsperb*^+/+^ males, respectively. (B and D) Sperm hyperactivation (% Motile) before (0 min) and after (90 min) incubation under capacitation conditions. (E and F) Flagellar waveform analysis of cauda spermatozoa. Tail motions of head tethered spermatozoa in the imaging chamber were recorded at 200 fps before (top, 0 min) and after (bottom, 90 min) capacitation. Overlays of flagellar waveforms for two beat cycles are color coded in time. (G and I) Beat frequency of spermatozoa before (0 min) and after (90 min) capacitation. Dots indicate α-angle of individual spermatozoa in each group. (H and J) Maximum angles of primary curvature in the midpiece (α-angle) of cauda sperm before (0 min) and after (90 min) induction of capacitation. Dots indicate the measured α-angle of individual spermatozoa in each group. Significance between groups was presented as *p<0.05, **p<0.01, ***p<0.001, and ****p<0.0001. *See also* Supplemental Figure 5 and 6.

### CATSPERδ ECD truncation phenocopies disrupting CATSPERβ-δ interface

To test the functional importance of the CATSPERβ–δ interaction reciprocally, we generated a transgenic mouse line expressing CATSPERδ lacking its canopy roof containing the CATSPERβ–δ binding region (**Supplemental Fig. 7A-C**). This line expresses only the canopy pole of CATSPERδ and therefore lacks the β–δ binding region; it was crossed onto a *Catsperd*-knockout background (*Catsperd*^−/−;TG+^).

*Catsperd*^−/−;TG+^ males were infertile (**Fig. 5A**), and CatSper proteins were undetectable in their spermatozoa although expressed in testes (**Fig. 5B**; **Supplemental Fig. 7D-F**). Testicular CATSPER1 protein levels were reduced in *Catsperd*^−/−;TG+^ mice (**Fig. 5C**), mirroring observations in *Catsperb*^Δ/Δ^ and *Catsperb*^−/−^ mice and aligning with trends reported in other CatSper canopy subunit-knockout models [29, 30]. Functionally, spermatozoa from both genotypes failed to undergo hyperactivation, displaying lower velocities and reduced beat amplitudes relative to wild type (**Fig. 5D-G**). Also, after the capacitation, P-Tyr was enhanced in *Catsperd*^−/−^ and *Catsperd*^−/−;TG+^ sperm than wildtype (**Supplemental Fig. 7G**). Accordingly, whereas wild-type sperm normally switch from circular to more linear trajectories under free-swim conditions in viscous media (0.5% methylcellulose) after capacitation [32, 51], both *Catsperb*^−/−^ and *Catsperd*^−/−;TG+^ spermatozoa swam poorly and failed to exhibit the efficient, linear movement at high viscosity (**Fig. 5H**). Consistent with these phenotypes, AlphaFold-Multimer predictions indicated that truncated CATSPERδ protein loses the ability to bind CATSPERβ and CATSPERε, independent of tag configuration (**Supplemental Fig. 8**). Together, these results demonstrate that an intact CATSPERβ–δ interaction within the canopy is essential for CatSper channel assembly, sperm function, and male fertility.

**Figure 5.**
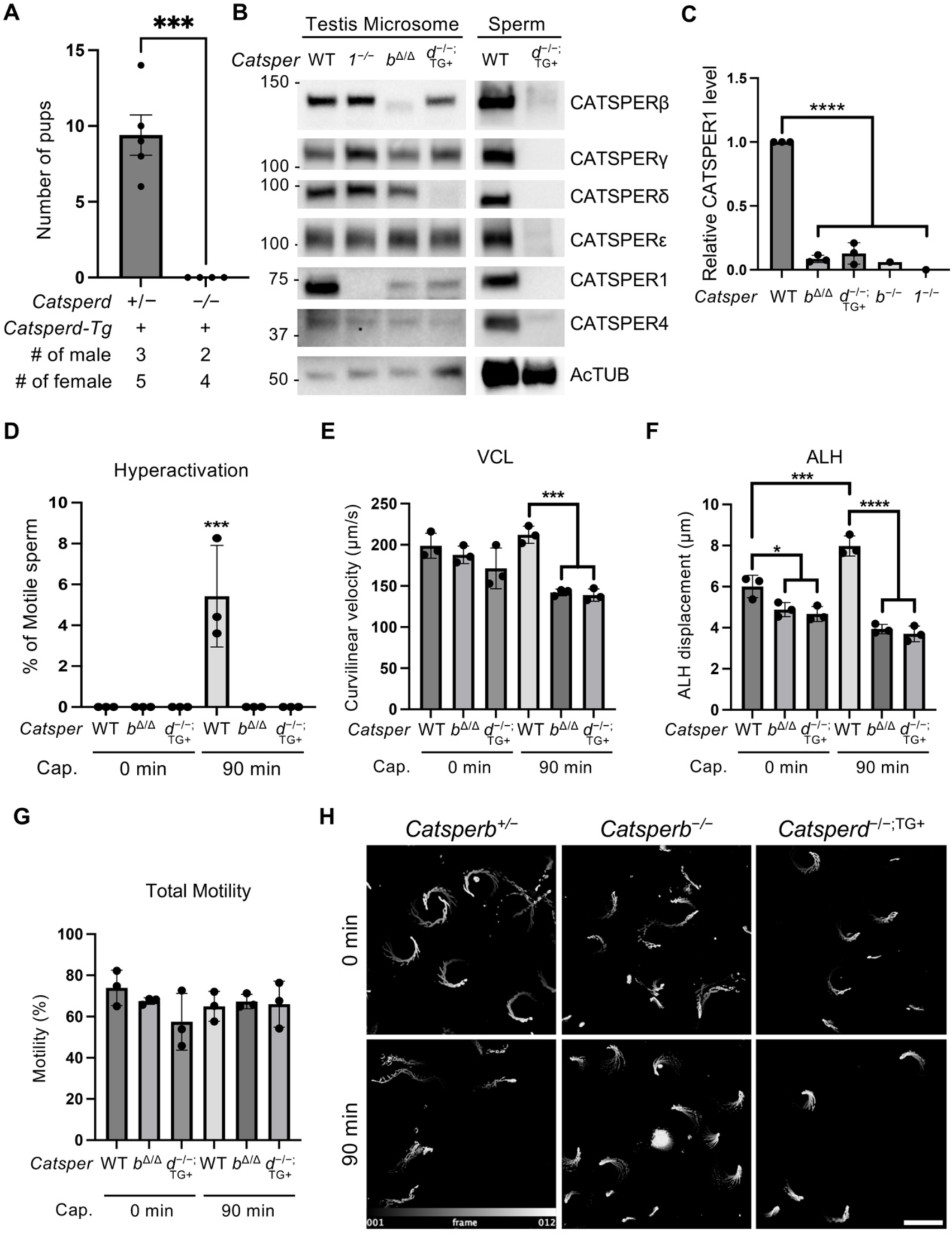
*Catsperd*^−/−;TG+^-transgenic male shows phenocopy of *Catsperb*^Δ/Δ^. (A) Percentage pregnancy rates of fertile females mated with transgenic males. (B) Immunoblot of testis microsome lysate and sperm from wildtype, *Catsper1*^−/−^, *Catsperb*^Δ/Δ^, and *Catsperd*^−/−;TG+^. (C) Quantification of CATSPER1 in different phenotypes. (D-G) Motility and correlative indicators of sperm wildtype, *Catsperb*^Δ/Δ^, and *Catsperd*^−/−;TG+^ before (0 min) and after (90 min) induction of capacitation in vitro. (D) Ratio of hyperactivated spermatozoa in motile spermatozoa before and after capacitation. (G) Motility of sperm before and after capacitation. (E) Curvilinear velocity of motile spermatozoa before and after capacitation. (F) Amplitude of lateral head before and after capacitation. (H) High-viscosity test of *Catsperd*^−/−;TG+^ sperm. Significance between groups was presented as *p<0.05, ***p<0.001, and ****p<0.0001. *See also* Supplemental Figure 7 and 8.

To probe the molecular basis of this interaction, we next tested whether predicted hydrogen-bonding residues contribute to CATSPERβ–δ binding. Five key CATSPERβ residues predicted to form hydrogen bonds with CATSPERδ were individually substituted with alanine (**Fig. 6A**) and CATSPERδ was co-expressed in HEK293T cells with either wild-type or mutant CATSPERβ. Co-immunoprecipitation confirmed that both wild-type and alanine-substituted CATSPERβ retained detectable interaction with CATSPERδ (**Fig. 6B**). However, the alanine-substituted mutant CATSPERβ exhibited a trend of reduced binding affinity for CATSPERδ mutants compared to the wild-type CATSPERβ (**Fig. 6C**), indicating that hydrogen bonding at the β–δ interface helps stabilize the interaction, especially considering full-length CATSPERβ and CATSPERδ protein are tested for the association.

**Figure 6.**
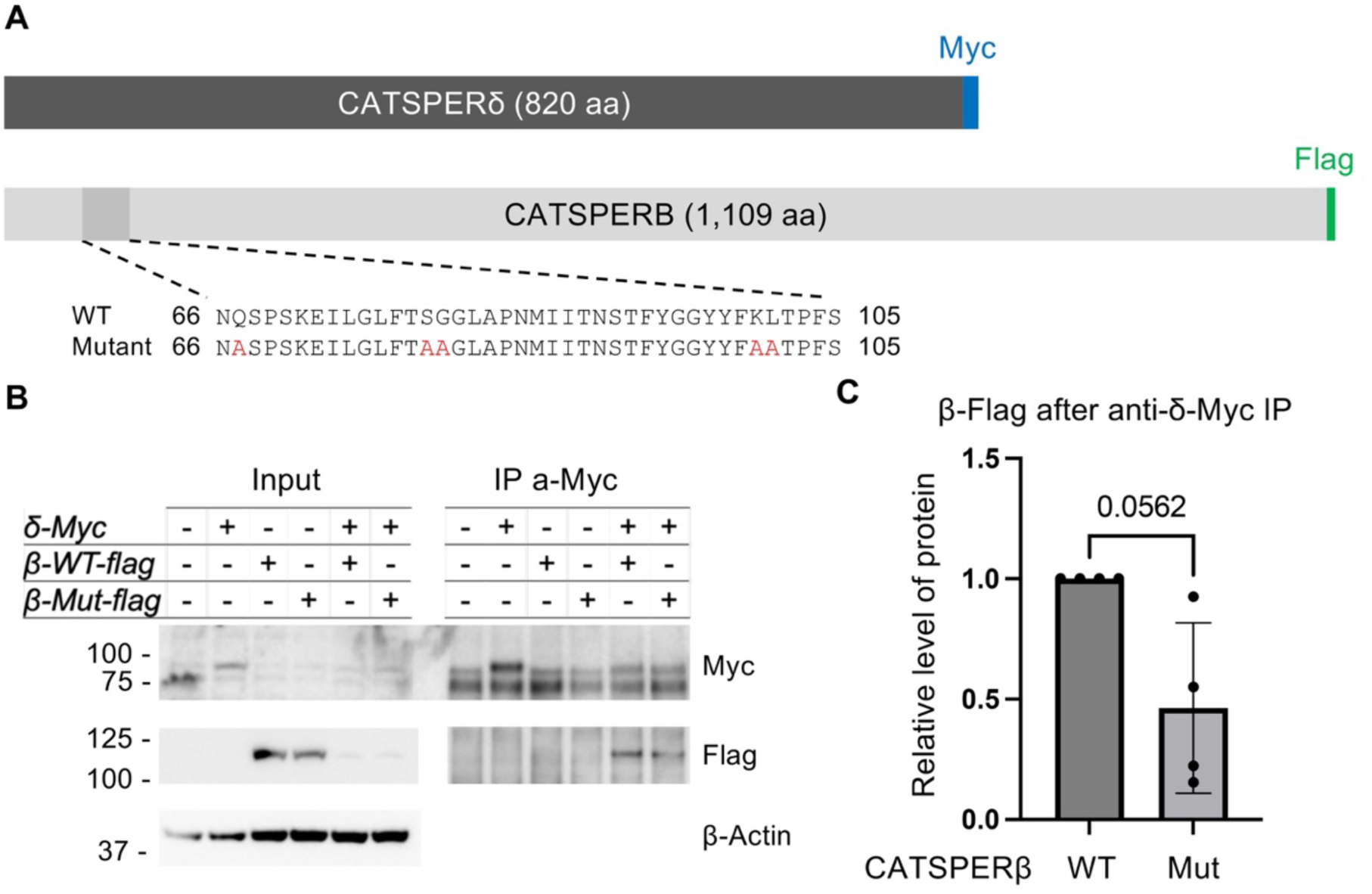
Reduced binding affinity of mutant CATSPERβ to CATSPERδ in a heterologous expression system. (A) Schematic of tagged constructs used for the binding assay. A Myc tag and a Flag tag was appended to the C-terminus of CATSPERδ and CATSPERβ, respectively. Five residues within the predicted CATSPERδ-binding interface of CATSPERβ were substituted with alanine to generate the mutant CATSPERβ construct. (B) Co-immunoprecipitation and immunoblot analysis of tagged proteins following anti-Myc immunoprecipitation from HEK293T cells co-expressing CATSPERδ-Myc with either wild-type or mutant CATSPERβ-Flag. (C) Quantification of the relative levels of wild-type and mutant FLAG-CATSPERβ co-immunoprecipitated with MYC-CATSPERδ. Values are normalized to wild-type (set to 1) within each independent experiment (n=4). Data represent mean ± SEM.

## Discussion

### The extracellular canopy serves as an obligate scaffold for hierarchical CatSper channel assembly

In mice, genetic abrogation of the pore-forming subunits does not measurably reduce the testicular protein levels of other pore subunits or auxiliary canopy components, suggesting that the core channel is not required to maintain canopy stability during spermatogenesis [29]. In contrast, loss of canopy subunits destabilizes the pore. For example, *Catsperd* or *Catspere* knockout lines exhibit reduced testicular CATSPER1 levels while leaving other canopy subunits largely intact [29, 30]. Our current findings significantly extend this hierarchy by demonstrating that CatSper holo-complex assembly is sensitive not only to the complete loss of an individual canopy subunit, but also to the precise disruption of a localized intra-canopy protein-protein interaction (PPI) interface.

Here, structural modeling and biochemical studies converge to establish CATSPERβ-δ interface as an indispensable organizing center for canopy assembly. AlphaFold-Multimer predictions indicate that an in-frame deletion of the exon 4-encoded 49 amino acid residues within CATSPERβ reduces intermolecular proximity at this structural junction. Alanine substitution of five hydrogen-bonding residues at this predicted interface showed a trend of reduced binding affinity *in vitro*. Co-IP experiments in *Catsperb*^Δ/Δ^ testes confirmed that this internally deleted variant fails to associate with CATSPERδ and is excluded from the broader CATSPERγ immunocomplex, proving that this local interface governs overall canopy roof integrity. This structural requirement is reciprocally corroborated by our *Catsperd*^−/−;TG+^ transgenic line; expressing an ECD-truncated CATSPERδ that lacks the β-binding domain faithfully phenocopies the fertility defects and total CatSper absence seen in the *Catsperb* mutant.

Our data support a model in which extracellular canopy scaffolds in a specific order and supports the integrity of pore-forming subunits that would subsequently assemble (**Supplemental Fig. 9**). This sequential dependency aligns perfectly with the temporal expression profile during spermatogenesis, where transcripts encoding the single TM canopy subunits with large ECDs are expressed prior to the pore-forming α-subunits [21, 32, 33]. In the mature sperm, the canopy is also suggested to regulate the channel activity (ref – Hwang 2026) rather than serving as a passive structural enclosure.

Although *Catsperg* knockout phenotype has not yet been reported, this hierarchical assembly model and its interdependent protein levels on other canopy subunit-knockout in mature sperm allows us to predict that genetic loss of CATSPERγ would similarly compromise CatSper biogenesis. This hypothesis is strongly supported by its obligate structural position within the heterotetrameric canopy and its contribution to extracellular interactions between dimers (“inter”-dimeric interfaces) in the zigzag array. Investigating whether a *Catsperg* deficiency phenocopies the complete structural collapse observed in our CATSPERβ-δ models remains a future avenue for validating the universality of this canopy-first assembly line.

### Species-specific variances in subunit persistence suggest distinct quality control mechanisms in mouse and human sperm

While our murine models demonstrate an absolute interdependence where sub-complex intermediates, if incomplete, are targeted for degradation or retained in cell body during spermatogenesis [29, 30], comparative evidence suggest intriguing species-specific differences in channel assembly and quality control. In human sperm, this divergence is clearly highlighted by a patient carrying a novel copy number variation (CNV) in *CATSPER2* [52]. Despite a profound reduction in CATSPER2 expression that abolished all measurable CatSper currents, the remaining pore-forming subunits (CATSPER1, CATSPER3, and CATSPER4) remained biochemically detectable in the patient sperm. This matches observations from distinct human cohorts with *CATSPER2* deletions, where CATSPERβ was reported to be absent [53], yet CATSPER3 and CATSPER4 were preserved and maintained their stereotypic quadrilinear tracking along the principal piece [54]. Together, these clinical observations suggest that the stringency of ER quality control, sub-complex stability, or trafficking checkpoint mechanisms may diverge significantly between mouse and human germ cells during spermatogenesis.

Given these variations, it will be informative for future diagnostic studies to evaluate the presence and localization of residual canopy and pore subunits in sperm from infertile patients harboring in-frame indels and/or missense *CATSPERE* mutations that abolish overall CatSper function [39, 40, 55]. In patients with an in-frame deletion, the loss of two amino acids in the *CATSPERE* stem leads to an absence of functional CatSper channel in sperm. Thus, structural completeness is important for forming a functional channel during spermatogenesis. Fully dissecting these species-specific biogenesis hierarchies will be critical for targeting mouse models into accurate diagnostic tools for human idiopathic male infertility. However, biochemical and high-resolution structural studies using human sperm samples remain constrained by sample availability. Expanding large-scale clinical biobanking efforts and performing rigorous, quantitative biochemical profiling of diverse human pathogenic variants will be essential future directions.

### The structural position of CATSPERβ establishes a key interaction hub for CatSper biogenesis and higher-order organization

Cryo-EM reconstructions reveal that the heterotetrameric canopy lies immediately above the tetrameric pore-forming channel (PDB: 7EEB). Within a single monomeric holo-complex, CATSPERβ engages adjacent δ and γ subunits extracellularly at the canopy roof and while anchoring to CATSPER4 via its single TM domain (**Fig. 1B**) (22). Beyond the individual holo-complex, CATSPERβ is an essential driver of the supramolecular zigzag arrays along the flagellar membrane (**Fig. 1A**) [23, 36]. Two CATSPERβ subunits form an intra-dimer β-β interface that links the canopy across a dimer [23, 36]. Recent structural work suggests that these higher-order longitudinal rows are further stabilized by intermolecular disulfide bonds at both the intra-dimer β-β interface and the inter-dimeric γ-γ interface [36]. Because the relevant cysteine residues are highly conserved among internal fertilizers [36], these covalent cross-links likely contribute to conformational stabilization or dynamic remodeling of the signaling nanodomains. This redox-dependent stabilization is intriguing given the stark physiological transition sperm must navigate, moving from a highly reducing environment in the testes and epididymis into the significantly more oxidizing milieu of the female reproductive tract [56].

At the lipid bilayer, CATSPERβ additionally interacts with the small auxiliary subunits CATSPERθ and CATSPERη at the intra-dimer interface [16, 30, 35]. Genetic loss of CATSPERθ does not broadly reduce pore or canopy protein levels in the testis, yet it selectively prevents the stable incorporation of CATSPER4 into the channel complex [35]. Taken together, the multi-layered connectivity of CATSPERβ—spanning from the extracellular canopy to the transmembrane interface dimeric interface—distinguishes it as the premier architectural linchpin of the CatSpermasome.

Consequently, targeting CATSPERβ-dependent interaction nodes presents an exceptionally promising avenue for non-hormonal male contraceptive development. Small-molecule therapeutics or peptidomimetics engineered to selectively occupy the CATSPERβ-δ interface and/or targeted protein degradation approaches such as PROTAC or molecular glue degraders designed to interfere with CATSPERβ-δ interaction during spermatogenesis would disrupt the primary scaffolding step of channel biogenesis with exquisite specificity. By leveraging AI-driven molecular docking and large-scale screens to target this discrete, sperm-specific interaction hub, it may be possible to arrest channel assembly long before the complex can reach the flagellar membrane, effectively blocking hyperactivated motility and fertilization.

## Author Contributions

Author contributions: J.-J.C. conceptualized the study. J.-N.O. developed the methodology. J.-N.O., H.W., T.W., and X.H. performed the investigation. J.-N.O. and H.W. visualized the data. J.-N.O. and H.W. conducted the formal analysis. J.-N.O., H.W., and J.-J.C. validated the work. J.-N.O. and H.W. curated the data. L.K.K. and J.-J.C. provided resources and J.J.C. supervised the project. J.-N.O. and J.-J.C. wrote the original draft and revised and edited the manuscript with the feedback from all authors.

## Acknowledgment

We thank M. Ikawa at Osaka University for sharing the pClgn-mCherry vector, J.Y. Hwang for generating *Catsperd*-*Tg* construct, R. Dai Pra for his support and initial characterization of *Catsperb* founder mice. J.-N. Oh was a recipient of a Lalor postdoctoral fellowship. J.-J.Chung is a Francis G. Kingsley Fellow at Yale School of Medicine.

## Movie Caption

**Movie S1. Comparison of atomic model and AlphaFold-multimer predicted CatSper canopy.** (A) Rotational movie of CATSPERβ, γ, δ, and ε that comprise the canopy of the CatSper complex (PDB: 7EEB). (B) the canopy at 70% transparency. (C) Fitting of AlphaFold-multimer predicted whole CATSPERβ-δ and CATSPERβ-γ ECDs. δ (green), γ (salmon), and β (light blue).

**Movie S2. Impact of the deletion within CATSPERβ NTD on β-δ and β-γ interactions predicted by AlphaFold multimer.** Lateral rotation movie showing the changes in the interfaces. CATSPERβ (light blue), CATSPERδ (green), and CATSPERγ (salmon).

**Movie S3. Vertical rotation of the CATSPERβ-δ AlphaFold multimer structure.** (A) CATSPERδ (green, 70% transparency), CATSPERβ (light blue), and CATSPERβ mutant (blue). (B and C) CATSPERδ (green, 70% transparency) and overlayed CATSPERβ (wildtype and mutant). The exon 4 encoded region in CATSPERβ is highlighted in magenta. Wildtype CATSPERβ has 70% transparency (C).

**Movie S4. Superimposed ECD structures of wildtype and mutant CATSPERβ.** (A) Structure alignment of CATSPERβ (light blue) and CATSPERβ mutant (blue). (B and C) Structure alignment of CATSPERβ (light blue, 70% transparency) and CATSPERβ mutant (blue). The peptide sequence corresponding to exon 4 in CATSPERβ is marked in magenta.

**Movie S5. Motility of tethered *Catsperb-*knockout and in-frame deletion mutant spermatozoa before (0 min) and after (90 min) induction of capacitation.** (A) *Catsperb*^+/−^ and *Catsperb*^−/−^. (B) *Catsperb*^+/+^ and *Catsperb*^Δ/Δ^.

