## Supplementary material for "CATSPERβ–δ Interaction Governs Hierarchical CatSper Holo-complex Assembly and is Essential for Male Fertility": Oh et al_Supplemental

**Supplemental Fig.1-Supplemental Fig. 9**
**Movie S1-S5**

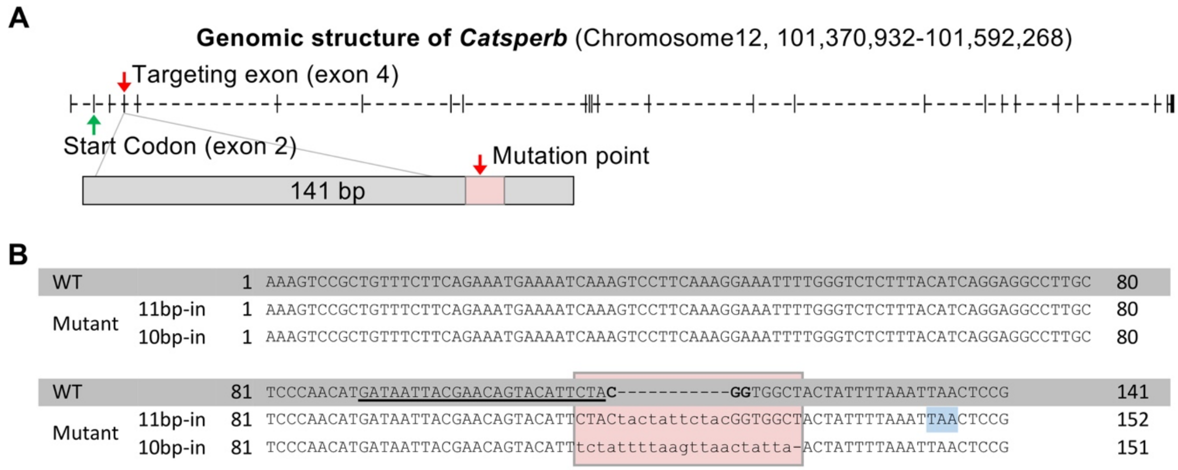


**Supplementary Figure 1. Genomic structure of WT and mutant *Catsperb* alleles and edited transcripts**, **related to Figure 1**. (A) Genomic structure of *Catsperb* gene and schematic of exon 4 editing targeted by CRISPR/Cas9. The red box indicates the mutated region in the mouse lines. (B) Exon 4 sequence of *Catsperb* transcripts from testes of wild-type and mutant animals (11-bp insertion: knockout; 10-bp insertion: truncated). The gRNA sequence is underlined, and the PAM sequence is in bold. The blue box indicates the stop codon in the 11-bp insertion (knockout) allele.

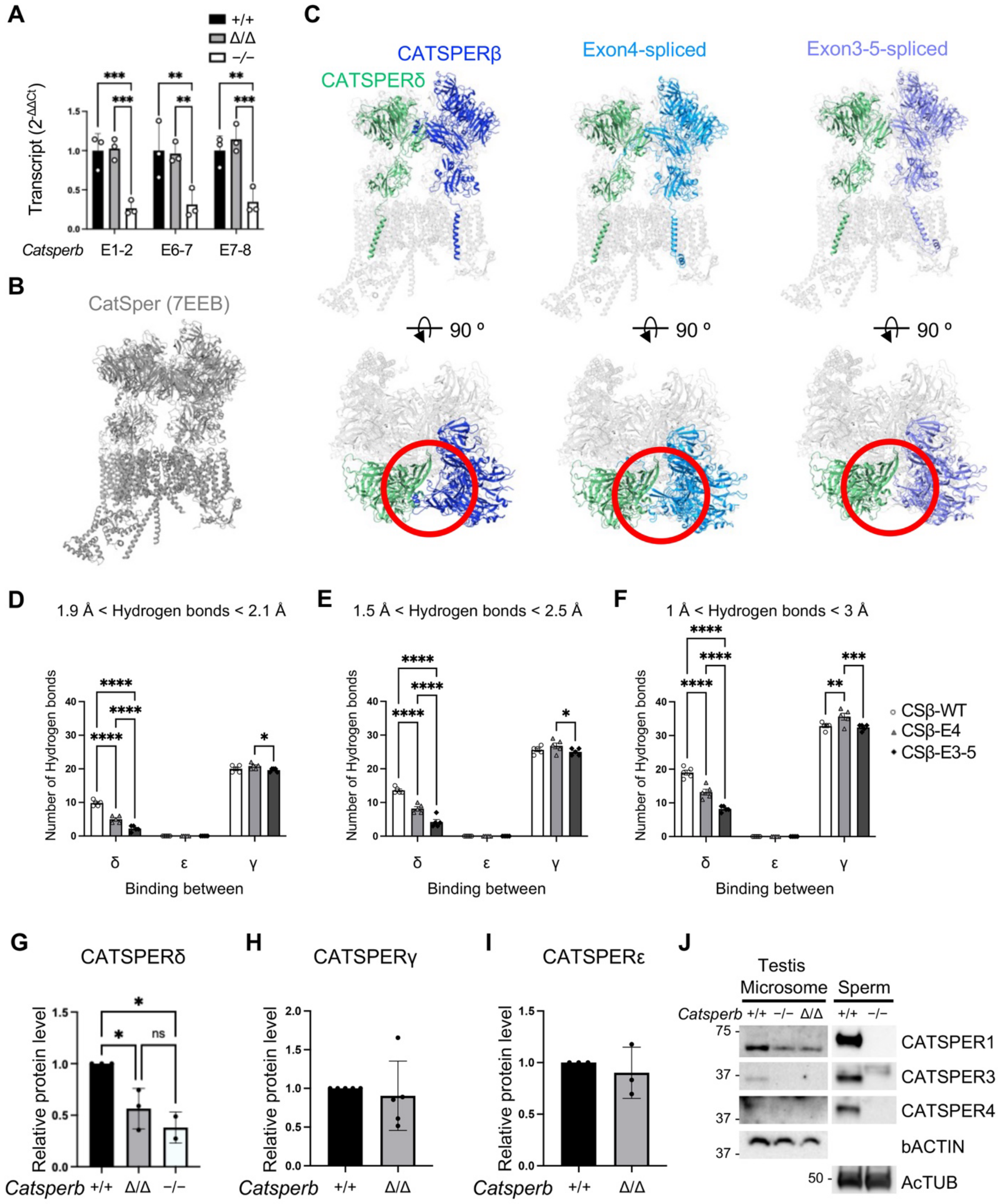


**Supplementary Figure 2. Predicted effects of CATSPERβ NTD truncation on the CATSPERβ-CATSPERδ interaction by AlphaFold, related to Figure 2.** (A) Relative level of *Catsperb* transcripts in adult testes. Transcripts levels were quantified using intron-flanking primers targeting exon 1-2, 6-7, and 7-8 (the exon 1-2 result is shown in Figure 2F as a representative). (B) Atomic model of the CatSper channel complex (PDB: 7EEB). (C) Fitting of predicted β mutant models into the CatSper complex. Predicted structures of CATSPERδ and CATSPERβ (left), CATSPERδ and CATSPERβ-Mutant (exon 4-spliced; center), and CATSPERδ with CATSPERβ-Mutant (exon 3, 4, 5-spliced; right) are shown. Bottom panels show the CATSPER δ and β interface. CatSper components other than CATSPER δ and β are shown at 50% transparency. (D–F) Number of hydrogen bonds between CatSper canopy proteins and CATSPERβ-E4 or -E3-5 (exon 4 or exon 3-5-spliced) at different distance thresholds: 0.1 Å (D), 0.5 Å (E), and 1 Å (F). (G-I) Quantification CATSPERδ (G), CATSPERγ (H), and CATSPERε (I) in testis microsome of wildtype, *Catsperb*$^{\Delta/\Delta}$, and *Catsperb*$^{-/-}$. (J) Immunoblot analysis of CatSper pore subunit protein levels in the solubilized testis microsomes from wild-type, *Catsperb*$^{-/-}$, and *Catsperb*$^{\Delta/\Delta}$ mice.

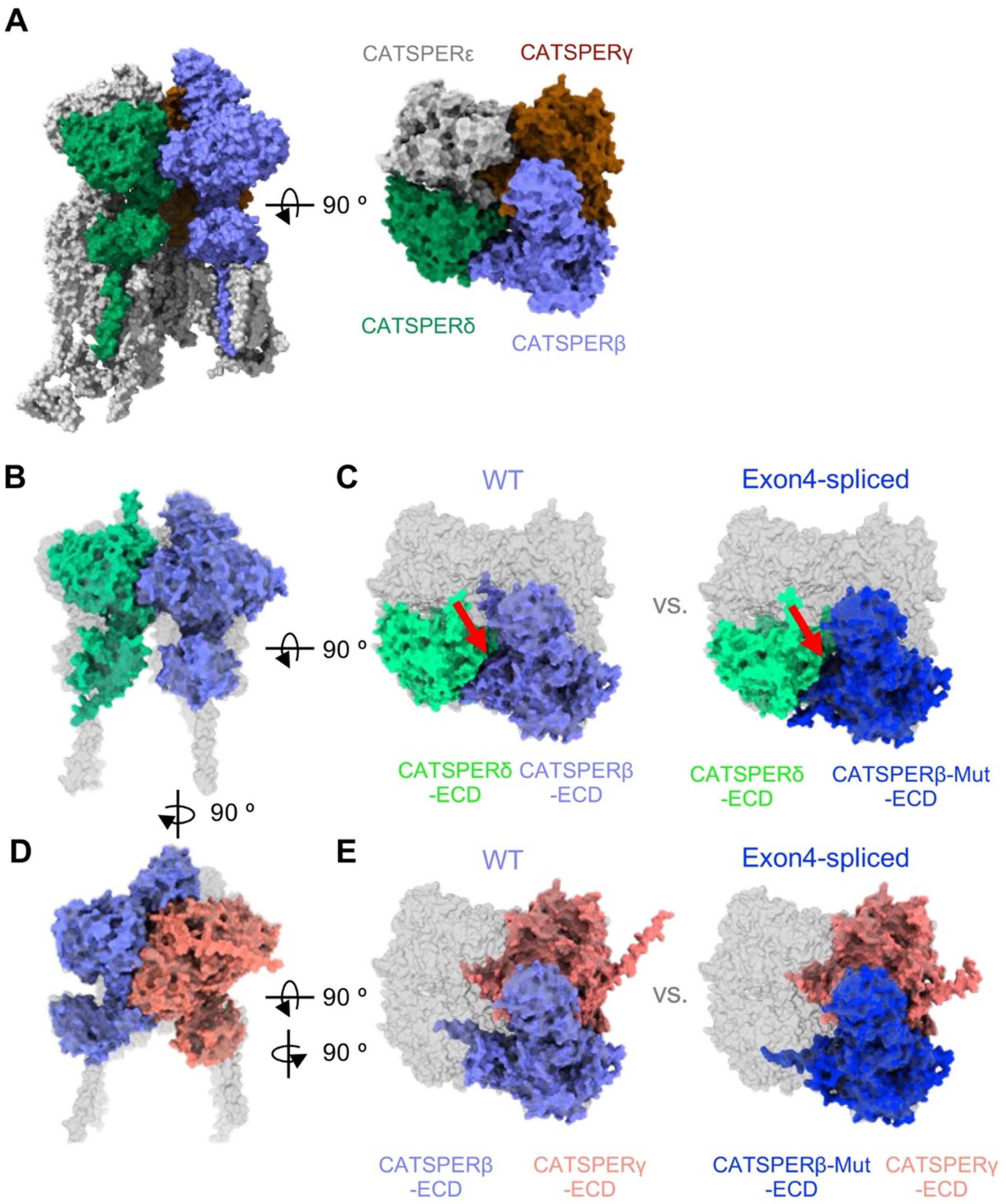


**Supplementary Figure 3. Canopy structure in the CatSper complex and fitting of β-γ and β-δ ECD multimer models from AlphaFold predictions, related to Figure 2.** (A) Canopy region comprising CatSper subunits (CATSPERβ, γ, δ, and ε) highlighted within the CatSper complex (PDB: 7EEB). *Left*, side view; *Right*, top-down view. (B-E) Fitting of multimer predictions into the canopy structure of the CatSper complex (70% transparency). CATSPER δ-CATSPERβ ECDs (B) and comparison with CATSPERδ- truncated CATSPERβ mutant ECDs (C). CATSPERγ-CATSPERβ ECDs (D) and comparison with CATSPERγ-truncated CATSPERβ mutant ECDs (E). Color key: δ (green), β (light blue), β-mutant (blue), γ (salmon). *See also* Movie S1.

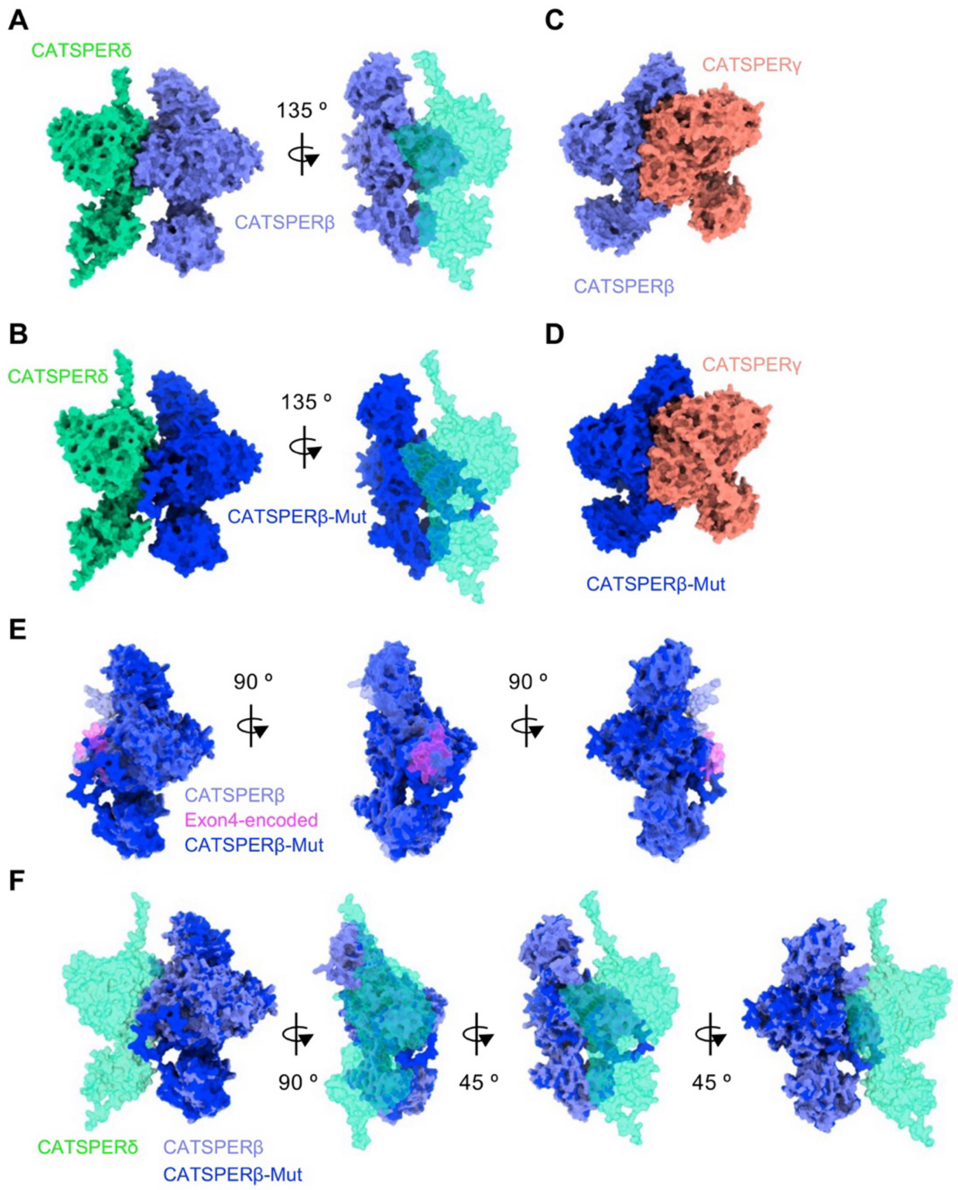


**Supplementary Figure 4. Predicted protein-protein interactions among extracellular domains in the CatSper canopy, related to Figure 2.** (A and B) AlphaFold-Multimer predictions of CATSPERδ-CATSPERβ (A) and CATSPERδ-CATSPERβ mutant (B) interactions. (C and D) AlphaFold-Multimer prediction of CATSPERγ-CATSPERβ (C) and CATSPERγ-CATSPERβ mutant (D) interaction. (E) Structural alignment of CATSPERβ and the CATSPERβ mutant. The peptide sequence encoded by exon 4 in CATSPERβ is shown in magenta. (F) Multimer structure predictions of CATSPERδ-CATSPERβ and CATSPERδ-CATSPERβ mutant interactions. Color key: δ (green), γ (orange), β (light blue), and truncated β (blue). Structures are shown at 70% Transparency. *See also* Movies S2, S3 and S4.

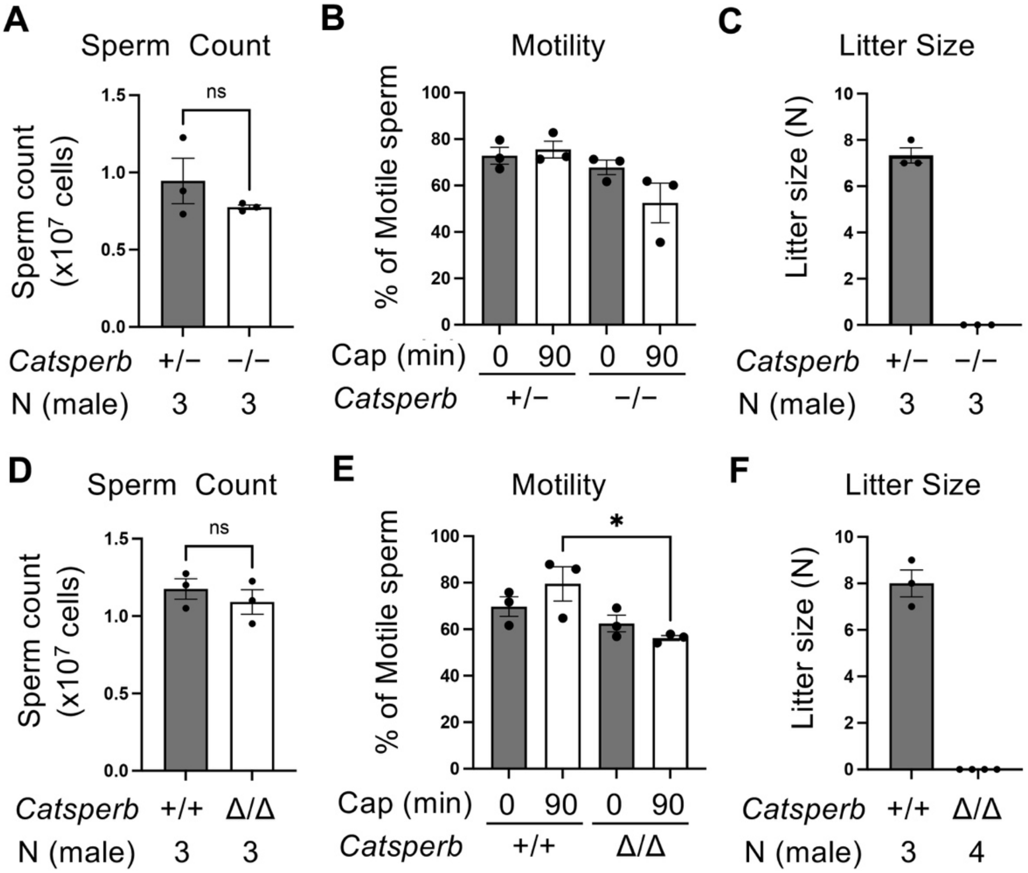


**Supplementary Figure 5. Complete knockout and truncation mutation of CATSPERβ impair male fertility without compromising spermatogenesis or overall sperm motility, related to Figure 4.** (A and D) Number of spermatozoa in each cauda epididymis from *Catsperb*$^{-/-}$ (A) and *Catsperb*$^{\Delta/\Delta}$ (D) males, compared with their littermate *Catsperb*$^{+/-}$ and *Catsperb*$^{+/+}$ males, respectively. (B and E) Cauda sperm motility before (0 min) and after (90 min) induction of capacitation. (C and F) Number of pups per litter after natural mating. Statistical significance is indicated as *p<0.05, **p<0.01, ***p<0.001, and ****p<0.0001.

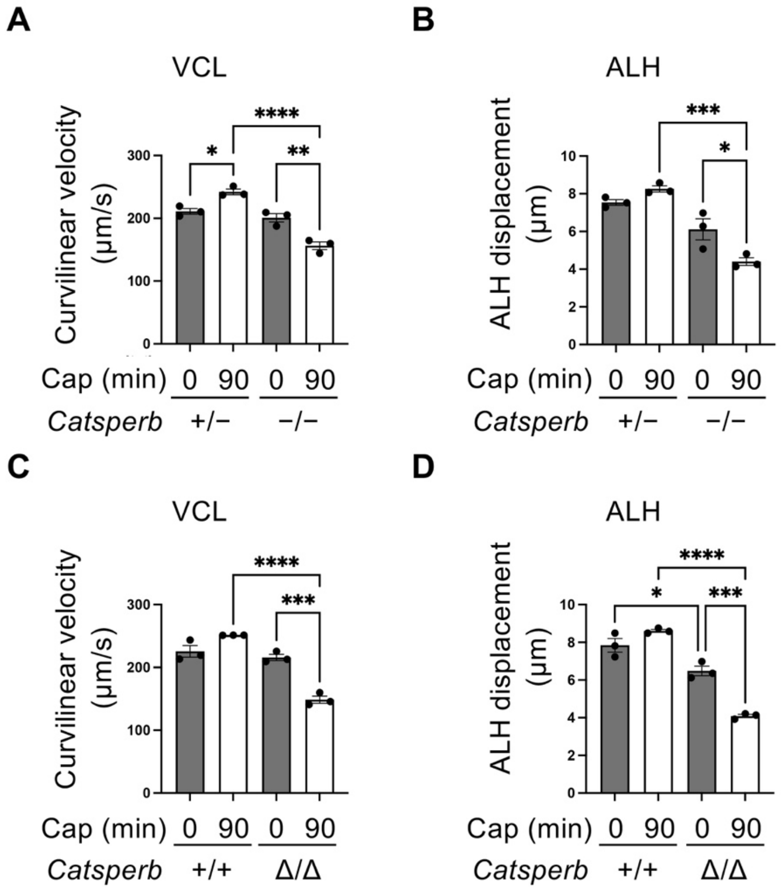


**Supplementary Figure 6. Complete knockout and truncation mutation of CATSPERβ impair male fertility by altering sperm motility, related to Figure 4.** (A and C) Curvilinear velocity and (B and D) lateral head displacement of motile spermatozoa before (0 min) and after (90 min) induction of capacitation in vitro. Genotypes compared were *Catsperb*$^{+/-}$ vs. *Catsperb*$^{-/-}$ (A and B) and *Catsperb*$^{+/+}$ vs. *Catsperb*$^{\Delta/\Delta}$ (C and D), respectively. Statistical significance is indicated as *p<0.05, **p<0.01, ***p<0.001, and ****p<0.0001.

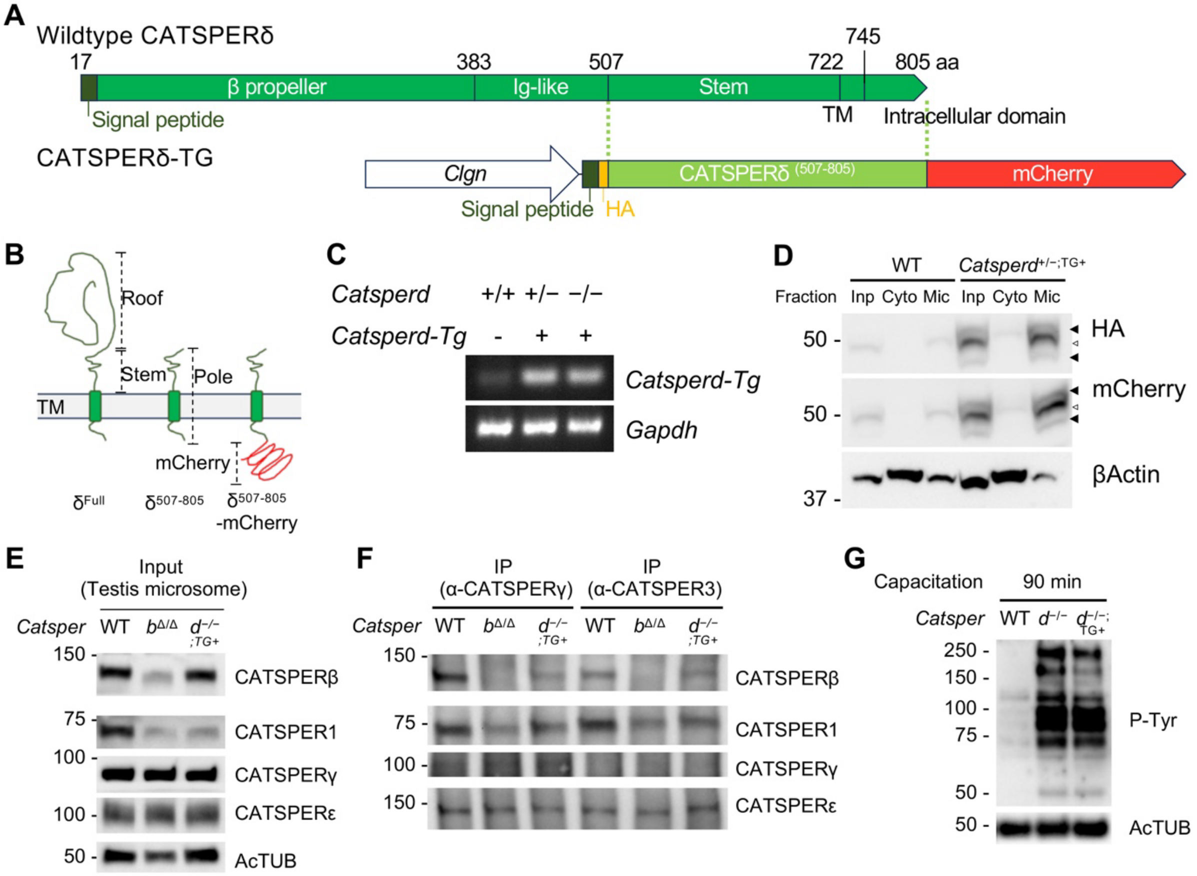


**Supplementary Figure 7. Generation of a mouse line expressing *Catsperd*-TG in the testis, related to Figure 5.** (A) Schematic diagram of CATSPERδ domain organization (UniProt: E9Q9F6) and the *Catsperd-Tg* transgene encoding an ECD-truncated CATSPERδ (CATSPERδ-TM) in mice. The transgene was fused to its native signal peptide, N-terminal HA and C-terminal mCherry tags, and driven by the *Clgn* promoter for testicular germ-cell-specific expression. (B) Predicted topologies of truncated (δ507-805) and full-length CATSPERδ proteins; the truncated construct retains only the canopy pole region. (C) PCR confirmation of transgene integration in the mouse line. (D) Expression of the transgene-encoded protein in mouse testis (HA-CATSPERδ-mCherry; CATSPERδ-TG) (Input; Inp, Cytosol; Cyto, Microsome; Mic). (E) Immunoblot analysis of CatSper subunit protein levels in the solubilized testis microsomes from wild-type, *Catsperb*$^{Δ/Δ}$, and *Catsperd*$^{-/-;TG+}$ mice. (F) Detection of CatSper proteins in anti-CATSPERγ and anti-CATSPER3 immunocomplexes. (G) Immunoblot of total capacitated sperm cell extract from wild-type, *Catsperd*$^{-/-}$, and *Catsperd*$^{-/-;TG+}$ mice by α-P-Tyr.

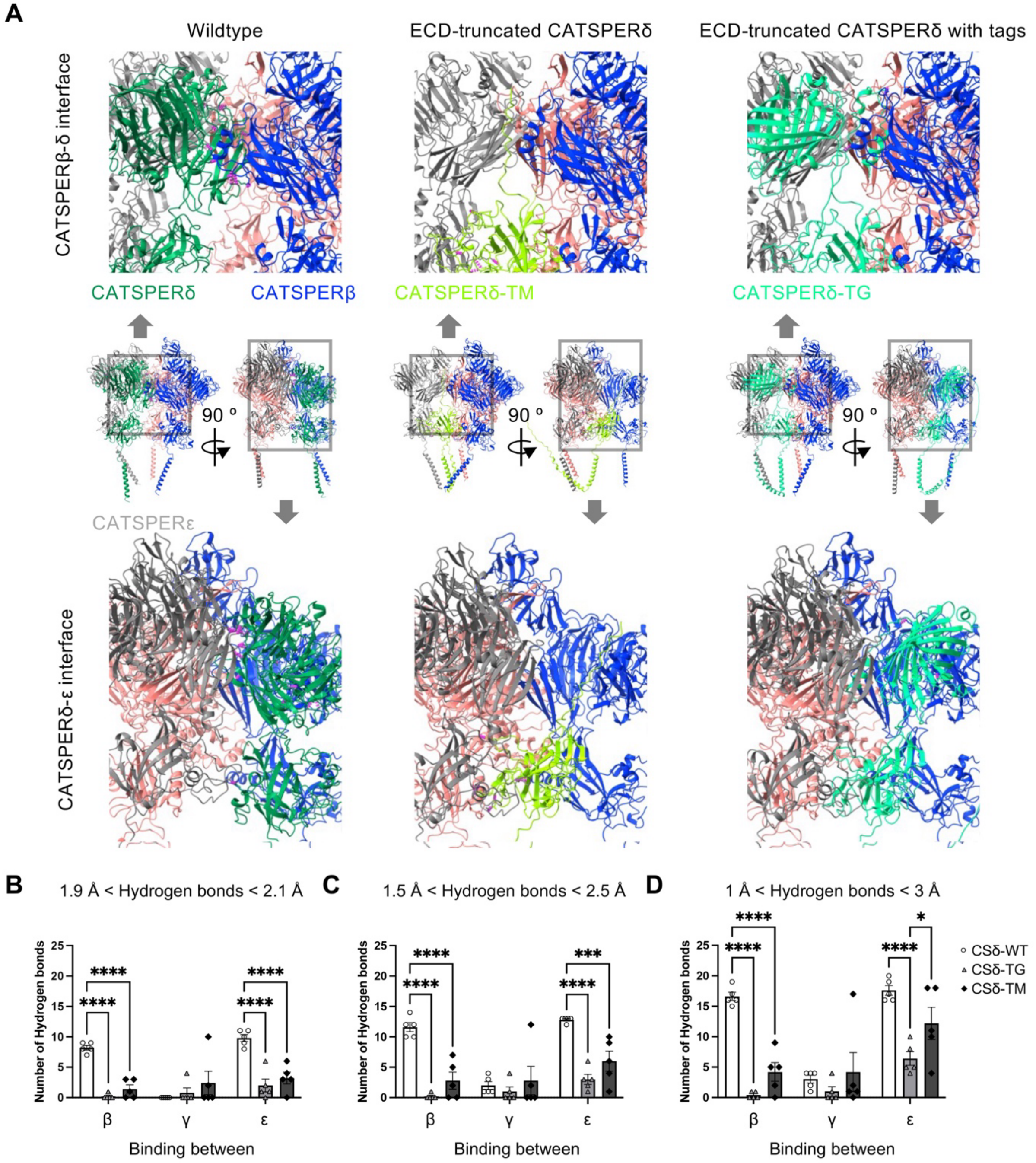


**Supplementary Figure 8. Prediction of hydrogen bonds from AlphaFold3 multimer structures, related to Figure 5.** (A) Interacting interfaces. Predicted hydrogen bonds between CatSper canopy proteins are depicted in dotted line (magenta). (B–D) Number of hydrogen bonds between CatSper canopy proteins and CATSPERδ-TM or -TG at different distance thresholds: 0.1 Å (B), 0.5 Å (C), and 1 Å (D).

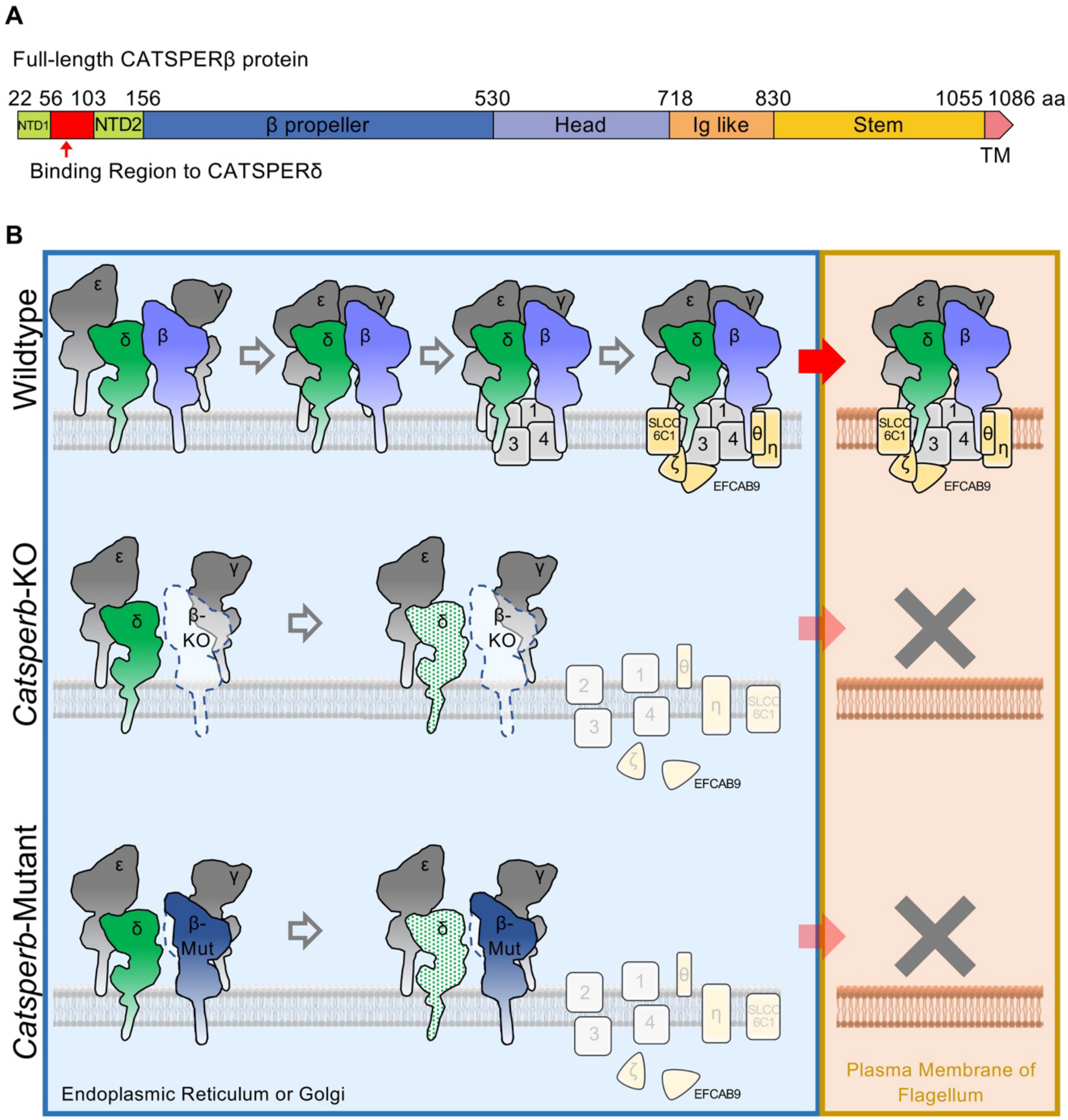


**Supplementary Figure 9. Proposed model of CatSper holo-complex assembly during spermatogenesis in WT, *Catsperb* knockout, and truncation mutant mice.** (A) Domain organization of CATSPERβ, with the region critical for protein-protein interaction with CATSPERδ highlighted in red. (B) Proposed assembly models of the CatSper channel complex in WT, *Catsperb* knockout, and truncation mutant mice. In WT testis, the four canopy subunits assemble into a tetrameric canopy beginning with CATSPERβ-CATSPERδ interaction; pore-forming subunits are then recruited to interact with each canopy stalk in pairs; likely followed by incorporation of the remaining subunits to form a functional CatSper holo-complex. In both *Catsperb*-KO and *Catsperb*-mutant testes, the loss of CATSPERβ-CATSPERδ interaction prevents proper tetrameric canopy assembly by destabilizing CATSPERδ and blocks all downstream steps in holo-complex biogenesis.

| | |
|---|---|
| CATSPERβ -ECD | VIHNKGKERTYFSCSGEGILTGLHTIKLFLTMDNLKVRCFFRNENQSPSKEILGLFTSGGLAPNMIITNSTFYGGYYFKLTPFSNRLEWLI<br>DIPRQNITVNTDIAAVEQWMIKITMHEGLNIYDTEGTLLDLVREPILQWNLGRVLTEMEVRDLYPEVNDIKVTKSPCANDVALIGFMMKPS<br>SNGVFIGKTISGFWTYKECIWHDLTEIIYAELKDEHQGLTVIDLVLTNHFLVILTSLGLYVSSDLRYPTTSQIKLSRAEFCGFERVDYIRG<br>NLWYNEKCFANRESFEVDYVTITFNRNRTLSESSSCFFSKEPFLHWLPCVFSTIKNEKSIPRVITFLIDQETDSGIYLFNVQDTKETYVTV<br>AMLKDGKPSPRPKFPSFHFPSTFTLPLGMIFHPRSHFLYVYGSQIWVSMDGGNTFEMLCNLFSHHVTKTSNSFYTSDIVFIVEDGRILTTK<br>AGLTTYSELGILKDAIFTLYYDQLGYIHKLTPENFDAGSKLLGHGNSGSIFGKRPDLGFEAILVPQYISTNEMYFFAHVPLTMPTNIQWKK<br>RFKTIHLGKTIEFSKTGLANIKNVYMHKTEPVGFQTSIHTEIIVPFGIENSKDSPCLLSDLEITYSGKLYYTIKLLSKNPLHELKSTDVEK<br>SVLIPGYSSFLIMNITDKWTASALATMPQAIKSNLKFLTGSWFLYNFGTAGGRKWSISTRQCNYWIQQDSLDFMSLNLVKYIDVGNTIDFQ<br>FKIIPKAMSTFPIPPVSMVVGNPGLVEVKTQGVFDLNENYYLDIHVSGRFFQKGSTSIALVLWEGSSKCYAITLLPTIKSSCSYLRTMHHT<br>PGRHIPPEDWISGVHKDSQGFNMIKTLPINYRPPSHMGISIPLTDNFYHADPSKPIPRNQFHKSKETGKYKQCANVTSRAMCNCSEHQKFS<br>HAVAFSDCKEKVHRFKFPVTQYPVVLEIFNERDKISAEPPYLVTMTEVNMRKNWQLKHNEPENVKKMKHYLEPLLKTPVYNPLGLNLTIQG<br>SELFHFKVSVVPGVSFCELSEEFQIYVDEVPLPFPG |
| CATSPERδ -ECD | HTLCRVHTVRTGKVFKSNIQLQGDPLFYAFPNTFVLKNVCKADISVYLGQKVFLTIDNFESSLLPLTVPKSLAVGVPSITSAHFVSGSLVL<br>FVISGKGYSYDYYENTWRKLEGISEPVSHISGDVCCFKGSFCLELSNNLFAYLRGGQIPGTNIYFSDNGGFSFQLMNTDKLSHLTGTLGGI<br>FHLHSMSQVGVLMVENNLGTFHYMEYPLNHSMGIAFSYKNLLEVIMKPYQRGFMVLWNQKSILVSSNSGQIVEHVRLIDQKIFTDLDVEHA<br>NINIYSVASNAYELAFLVAEDHLYYGSQSYMGTYVIKLPHQPLWSTHTSIYFEDIGILQVLTPVADPHFAAYDFDKCTVNVQSSLMDEKLA<br>LQPCNVELLESTMINTMFTIDMNSKLKLSALMIPRKGENPTPLVMVSNPHALGFKANLNEFGNTFDGNSKYKLDIELKQQHHWGNSDFNFT<br>ASIKRHAISSVTVDIADKTLSCVDLKPLSTLISVGCDMTKKIVVQNKISACTMGILNPVQLQKNYTYTIEKEAYDPINHNGEAQDDLIVFY<br>EYKDLGCPRLVYYDKPWKPVVELWKNGIVEEIMNAEYVISEINGLVTYSYSLTAATANCRSQPQNWSTFESDIENEEPFLWNRENYVSCHE<br>DNKDNPLLWPNVEYQVLGGQTNNKIIFGQRNGIYTFHLSVVDPYYSYCNLNTIFSVYVHGALPVTKFQPL |
| CATSPERγ -ECD | HCTWLLVLNKFEKVGLHLSKDRFQDHEPIDTVAKVFQKLTDSPIDPSENYLSFPYYLQINFSCPGQNIEELARKGHLMGMKPMVQINYMYS<br>VNFYRWEMENVQILMEAAPMRSTGYCPAEAMCVLNWYTPMPFKNGSVVSSVDIYTNGIGPFVSKKRFYVNMNGFLKRDASGKSLFAIGYES<br>LVLKSSHFRLSKSRPLWYTVNHAPVFILGGFYDEKSILFSDSNFQDYVLLELSIDSCWVGSFYCPILGFSATIHDAIATESTLFIRQNQLV<br>YYFTGTYITLFDKSHGSSRWVRVLPSECIKRLCPVYASGNGSEYVLALTTGKNEGYIHIGTITDGLVSFEMVPDGWSVCEKLPGKNCSIDW<br>ATYIADERNLLLLVKIDSGQFYLVNFNTEFKTLNILYKIPEFIPEAKELDFLVLLDTVTYTNTPMTPKGLFFNTLNNMLYIWGNFILQSYN<br>REEFIFLADFPKESTIKYMVNSFKGQMAVVTENEEIWYFLEGGYDVYQVVPSQGWETYHNLQKMQKSSFHSEDESLVSLFFEDGKLFQLVY<br>LFDVGKERLVKRLLPVGTLMEYNLPKPFTVVNQGNYQAISFTHTCPFKEIHLIDVPKKHHASRTESYVALPPLVSESLGFHNNNTLAVYQG<br>LVYYLLWLHSKYDKPYADPVHDPTWRWWQHKTKDKDYFFYLFSNRLAAEGIYINMNAYQKLYNMSGDYGIPDLFFLDKGNWFTITVVLLSH<br>QDTFTSSDSQGPTINVDKKLAIAVTIADPECLSVTVTQDVLLNRNAVINKIKVIDKKRCSEQGMIGRNIKKTSMMLKVLGAPGNCIQRTYL<br>GGIIQGFKVVPIFIGCPPGKRLAFDVSYTIMHSEEINKHYFDCVIKDAEMPCFLFRDLFQPFFLVQDLVTGDSGSFLGSYVLKVVGGGRTL<br>NTIRDYTEEEIFRYNSPLDTTNSLIWKTKVERTTEDKKFYIMSHESPGVEWLCLENSPCYDIIPQSIYPPEFFFKLLVSNRGVDNSTYCDY<br>KLTFIVHIHGLPL |
| CATSPERε -ECD | LWRYYINSQDYSIFSTRSSIKLEYEGNSFVSWKIPESCKVENTTSPKTTLHCKRAGIHTIKPIAGNQEVERHLTVDNSYICYLWYFTVVDV<br>YYNLSQIVTIWVYDPESASTEELIWTAKKPSLSSRVLTKQMNTLGQRPFIFTVEKRLTYHPGPLTSEGTWVIHLPMSSDDIAKVIRGNKVA<br>FQDCFIANLYFMLTYPMTIISEPPGYEPLTVPPGSPLMLSWDTCISTFALLATDQETFQTNDSFQTWTRVRAPPGILSDAQRHSLRDVIIF<br>DQGTLFLVDGTVYLRTEDEFTKLDESRGISETGILGFSKRRWCQIRYLYKLASKKSILIAWSKTTVYAGYATFRFVTLTDTAKLKDFLKLP<br>QTDTLEVMSVEYLWHPLEAAVLLSHCSVCTTNTRNIRIVIYSAIFQTWTLQDFELQLPKEAILEFRFLYSAMPDIIMWDQHHVYYSYKNFT<br>VVGTISTPSGETNLSSLSQGSKIHQVLTDRIGNVVVKMENNVMFYIKADITEAVILHTWVNTTAKTVVLFDKSFEVCILYYNENLDEKYQL<br>QTQPYPLILELQSINKDLGDWCPYLAFQHNIHSQFYHMDKGESLTIWSQIVYPENRGLYIVVEHYGSSVMTWTQNLEYEIASGFCTKTMIT<br>RFFQTTNYELVDNYYQLQKENTGLMLLQFRPSEFSRTCLTAKPVFEIDVGCDSSKYIMVRGFNKSRCQRRDFSYVIDKELLRESLSDNLKV<br>RYDVAKYGCPLTLELGQMFQPIVELYDENGFIKIVDANFILWEIHGRNDYTFNSTMEQNGCINEAQTWDSMIEENPDIPLDDVWGPQNYRP<br>CFSYAIGKPGDLGQPYEILNYSNKNHIKWPMTYAGMYVYRLKILDPNYSFCNLTTIFAIESLGMIPRS |
| Exon4-spliced CATSPERβ ECD | VIHNKGKERTYFSCSGEGILTGLHTIKLFLTMDNLFSNRLEWLIDIPRQNITVNTDIAAVEQWMIKITMHEGLNIYDTEGTLLDLVREPIL<br>QWNLGRVLTEMEVRDLYPEVNDIKVTKSPCANDVALIGFMMKPSSNGVFIGKTISGFWTYKECIWHDLTEIIYAELKDEHQGLTVIDLVLT<br>NHFLVILTSLGLYVSSDLRYPTTSQIKLSRAEFCGFERVDYIRGNLWYNEKCFANRESFEVDYVTITFNRNRTLSESSSCFFSKEPFLHWL<br>PCVFSTIKNEKSIPRVITFLIDQETDSGIYLFNVQDTKETYVTVAMLKDGKPSPRPKFPSFHFPSTFTLPLGMIFHPRSHFLYVYGSQIWV<br>SMDGGNTFEMLCNLFSHHVTKTSNSFYTSDIVFIVEDGRILTTKAGLTTYSELGILKDAIFTLYYDQLGYIHKLTPENFDAGSKLLGHGNS<br>GSIFGKRPDLGFEAILVPQYISTNEMYFFAHVPLTMPTNIQWKKRFKTIHLGKTIEFSKTGLANIKNVYMHKTEPVGFQTSIHTEIIVPFG<br>IENSKDSPCLLSDLEITYSGKLYYTIKLLSKNPLHELKSTDVEKSVLIPGYSSFLIMNITDKWTASALATMPQAIKSNLKFLTGSWFLYNF<br>GTAGGRKWSISTRQCNYWIQQDSLDFMSLNLVKYIDVGNTIDFQFKIIPKAMSTFPIPPVSMVVGNPGLVEVKTQGVFDLNENYYLDIHVS<br>GRFFQKGSTSIALVLWEGSSKCYAITLLPTIKSSCSYLRTMHHTPGRHIPPEDWISGVHKDSQGFNMIKTLPINYRPPSHMGISIPLTDNF<br>YHADPSKPIPRNQFHKSKETGKYKQCANVTSRAMCNCSEHQKFSHAVAFSDCKEKVHRFKFPVTQYPVVLEIFNERDKISAEPPYLVTMTE<br>VNMRKNWQLKHNEPENVKKMKHYLEPLLKTPVYNPLGLNLTIQGSELFHFKVSVVPGVSFCELSEEFQIYVDEVPLPFPG |
| Exon3-5-spliced CATSPERβ ECD | VIHNKDIAAVEQWMIKITMHEGLNIYDTEGTLLDLVREPILQWNLGRVLTEMEVRDLYPEVNDIKVTKSPCANDVALIGFMMKPSSNGVFI<br>GKTISGFWTYKECIWHDLTEIIYAELKDEHQGLTVIDLVLTNHFLVILTSLGLYVSSDLRYPTTSQIKLSRAEFCGFERVDYIRGNLWYNE<br>KCFANRESFEVDYVTITFNRNRTLSESSSCFFSKEPFLHWLPCVFSTIKNEKSIPRVITFLIDQETDSGIYLFNVQDTKETYVTVAMLKDG<br>KPSPRPKFPSFHFPSTFTLPLGMIFHPRSHFLYVYGSQIWVSMDGGNTFEMLCNLFSHHVTKTSNSFYTSDIVFIVEDGRILTTKAGLTTY<br>SELGILKDAIFTLYYDQLGYIHKLTPENFDAGSKLLGHGNSGSIFGKRPDLGFEAILVPQYISTNEMYFFAHVPLTMPTNIQWKKRFKTIH<br>LGKTIEFSKTGLANIKNVYMHKTEPVGFQTSIHTEIIVPFGIENSKDSPCLLSDLEITYSGKLYYTIKLLSKNPLHELKSTDVEKSVLIPG<br>YSSFLIMNITDKWTASALATMPQAIKSNLKFLTGSWFLYNFGTAGGRKWSISTRQCNYWIQQDSLDFMSLNLVKYIDVGNTIDFQFKIIPK<br>AMSTFPIPPVSMVVGNPGLVEVKTQGVFDLNENYYLDIHVSGRFFQKGSTSIALVLWEGSSKCYAITLLPTIKSSCSYLRTMHHTPGRHIP<br>PEDWISGVHKDSQGFNMIKTLPINYRPPSHMGISIPLTDNFYHADPSKPIPRNQFHKSKETGKYKQCANVTSRAMCNCSEHQKFSHAVAFS<br>DCKEKVHRFKFPVTQYPVVLEIFNERDKISAEPPYLVTMTEVNMRKNWQLKHNEPENVKKMKHYLEPLLKTPVYNPLGLNLTIQGSELFHF<br>KVSVVPGVSFCELSEEFQIYVDEVPLPFPG |

**Table S1. The protein sequences utilized for AlphaFold structure prediction**. Listed are the extracellular portion of CATSPER β, δ, γ, ε, and mutant CATSPER β sequences utilized for multimer structure prediction via AlphaFold. The sequences are sourced from the Protein Data Bank (PDB) entry 7EEB.

**Movie caption**

**Movie S1. Comparison of atomic model and AlphaFold-Multimer-predicted CatSper canopy.** (A) Rotational view of CATSPERβ, γ, δ, and ε that comprise the canopy of the CatSper complex (PDB: 7EEB). (B) Canopy shown at 70% transparency. (C) Fitting of AlphaFold-Multimer-predicted full extracellular domains (ECDs) for CATSPERβ-δ and CATSPERβ-γ. Color key: δ (green), γ (salmon), β (light blue).

**Movie S2. Impact of truncation in the CATSPERβ NTD on β-δ and β-γ interactions predicted by AlphaFold-Multimer.** Lateral rotation highlighting interface changes. Color key: CATSPERβ (light blue), CATSPERδ (green), CATSPERγ (salmon).

**Movie S3. Vertical rotation of the CATSPERβ-δ AlphaFold-Multimer structure.** (A) CATSPERδ (green, 70% transparency), CATSPERβ (light blue), and CATSPERβ mutant (blue). (B and C) CATSPERδ (green, 70% transparency) with overlaid CATSPERβ (wild-type and mutant). The exon 4-encoded region in CATSPERβ is highlighted in magenta; wild-type CATSPERβ is shown at 70% transparency in (C).

**Movie S4. Superimposed ECD structures of wild-type and mutant CATSPERβ.** (A) Structure alignment of CATSPERβ (light blue) and CATSPERβ mutant (blue). (B and C) Alignment with CATSPERβ shown70% transparency. The peptide sequence corresponding to exon 4 in CATSPERβ is marked in magenta.

**Movie S5. Motility of tethered *Catsperb* knockout and truncation-mutant spermatozoa before (0 min) and after (90 min) induction of capacitation.** Sperm from (A) *Catsperb*$^{+/-}$ and *Catsperb*$^{-/-}$ and (B) *Catsperb*$^{+/+}$ and *Catsperb*$^{\Delta/\Delta}$ males.
